# The Impact of a Natural Ingredients based Intervention Targeting the Nine Hallmarks of Aging on DNA methylation

**DOI:** 10.1101/2024.09.02.610828

**Authors:** Natalia Carreras-Gallo, Rita Dargham, Shealee P. Thorpe, Steve Warren, Tavis L. Mendez, Ryan Smith, Greg Macpherson, Varun B. Dwaraka

## Abstract

Aging interventions have progressed in recent years due to the growing curiosity about how lifestyle impacts longevity. This study assessed the effects of SRW Laboratories’ Cel System nutraceutical range on epigenetic methylation patterns, inflammation, physical performance, body composition, and epigenetic biomarkers of aging. A 1-year study was conducted with 51 individuals, collecting data at baseline, 3 months, 6 months, and 12 months. Participants were encouraged to walk 10 minutes and practice 5 minutes of mindfulness daily. Significant improvements in muscle strength, body function, and body composition metrics were observed. Epigenetic clock analysis showed a decrease in biological age with significant reductions in stem cell division rates. Immune cell subset analysis indicated significant changes, with increases in eosinophils and CD8T cells and decreases in B memory, CD4T memory, and T-regulatory cells. Predicted epigenetic biomarker proxies (EBPs) showed significant changes in retinol/TTHY, a regulator of cell growth, proliferation, and differentiation, and deoxycholic acid glucuronide levels, a metabolite of deoxycholic acid generated in the liver. Gene ontology analysis revealed significant CpG methylation changes in genes involved in critical biological processes related to aging, such as oxidative stress-induced premature senescence, pyrimidine deoxyribonucleotide metabolic process, TRAIL binding, hyaluronan biosynthetic process, neurotransmitter loading into synaptic vesicles, pore complex assembly, collagen biosynthetic process, protein phosphatase 2A binding activity, and activation of transcription factor binding. Our findings suggest that the Cel System supplement range may effectively reduce biological age and improve health metrics, warranting further investigation into its mechanistic pathways and long-term efficacy.

## INTRODUCTION

Aging is now known to be the single biggest risk for disease. In recent years, advances in aging interventions have been driven by increased interest in the effects of lifestyle on healthspan and lifespan. Genetic research [1], senolytic therapies [2], stem cell treatments [3], calorie restriction mimetics [4],[5], microbiome modulation [6], and epigenetic clock modulation [7] have collectively advanced the field of aging interventions. As medical treatments and therapies progress, aging, as the major contributor to illness and mortality, becomes increasingly prominent. Nutraceuticals are emerging as a promising tool to optimize health and prolong lifespan. The global dietary supplement market was estimated at USD 177.50 billion in 2023 and is expected to reach USD 192.65 billion in 2024 [8]. There are numerous health benefits behind certain supplements, such as folic acid and omega-3 fatty acids [9], but a nutraceutical that decreases biological age or slows the rate of aging has not been established.

Recently, the development of the Cel^123^ (Cel) by SRW Laboratories utilizes a three part formulation which is suggested to support functional areas and processes in the cellular system which typically decline with age. The Cel1 nutraceutical formulation contains 2-HOBA (hobamine^TM^), astragalus membranaceus extract (astragaloside), sophorae japonica extract (rutin), vitamin C, levomefolic acid, vitamin B12, zinc, and selenium. 2-HOBA supports DNA defense against free radical damage, as it neutralizes reactive carbonyl species that cause stress in cells [10]. This molecule binds to lipid free radicals, preventing them from binding to proteins and DNA, causing mutations that result in cellular damage [10]. Several studies support 2-HOBA’s ability to support healthy aging, DNA integrity, brain function, immune, and circulatory health [10]. Astragaloside is known to support telomere integrity and repair through stimulation of enzymes responsible for DNA repair [11]. Rutin strengthens the expression of antioxidant proteins and enzymes that protect against DNA damage associated with external stress, including UV exposure and other oxidative stress [12]. The antioxidant properties in vitamin C have shown therapeutic effects against oxidative stress, disorganization of chromatin, and telomere attrition, prolonging lifespan [13]. Deficiency in levomefolic acid, contributes to aging in elderly patients [14]. Vitamin B12, zinc, and selenium have been proven in several studies to contribute to improved immune function and decreased cellular aging through the reduction of oxidative stress, decreased DNA methylation, and telomere stability [15–18]. Vitamin B12, levomefolic acid, and zinc supplementation has additionally shown decreased epigenetic age measured by an epigenetic clock analysis calculated by the Horvath model [19], [20]. Selenium is considered an epigenetic modulator and correlates with at least one differentially methylated genome region [21].

The Cel2 supplement incorporates a proprietary blend of ingredients including nicotinamide mononucleotide (NMN), pterostilbene, astaxanthin, L-carnosine, vitamin D, and riboflavin. NMN is a systemic signaling molecule and NAD+precursor, boosting cellular levels, with downstream effects positively impacting insulin sensitivity, mitochondrial dysfunction, and extending lifespan [22]. Pterostilbene has demonstrated regulation of gene expression by altering epigenetic patterns [23]. Astaxanthin, a potent antioxidant, shields cell membranes from damage, promoting resilience against age-related oxidative stress [24]. L-carnosine exhibits versatile anti-aging effects by inhibiting glycation, scavenging free radicals, and supporting mitochondrial function, thereby enhancing cellular vitality [25]. Vitamin D modulates immune function and gene expression associated with cellular aging, bolstering resilience against age-related decline [26].

Riboflavin’s role as a cofactor in energy metabolism and antioxidant defense reinforces cellular health, mitigating age-associated oxidative stress [27].

The Cel3 formula contains apigenin, fisetin, oleuropein, EGCG, berberine, alpha lipoic acid, and withaferin A. Apigenin’s anti-inflammatory, CD38 inhibition and antioxidant properties protect cells from age-related stressors [28], while the natural senolytic compound fisetin modulates critical signaling pathways associated with the central nervous system linked to longevity [29]. Oleuropein, EGCG, and withaferin A contribute their antioxidant properties, bolstering cellular integrity, proteostasis and mTor inhibition [30], [31]. Berberine’s multifaceted impact on the AMPK pathway and downstream impact on cellular metabolism and inflammation further supports anti-aging mechanisms [32]. Alpha lipoic acid serves as both a mitochondrial cofactor and protects against oxidative damage [33].

Primary, integrative, and antagonistic sectors of the hallmarks of aging focus on the molecular, cellular, and systemic processes accounting for the morphological and functional decline that affects aging [34]. These hallmarks include DNA instability, telomere attrition, epigenetic alterations, loss of proteostasis, deregulated nutrient-sensing, mitochondrial dysfunction, cellular senescence, stem cell exhaustion, altered intercellular communication, disabled macroautophagy, chronic inflammation, and dysbiosis [34]. The Cel System supplement range was formulated to target pathways associated with the Hallmarks of Aging when combining Cel1, Cel 2, and Cel 3 formulas. Cel1 primarily targets DNA instability, telomere attrition, mitochondrial dysfunction, cellular senescence, and chronic inflammation. w Cel2 synergistically targets aging pathways relating to mitochondrial dysfunction, dysbiosis, stem cell health and intercellular communication Cel3 targets cellular senescence, proteostasis, and nutrient sensing and macroautophagy.

Epigenetic biomarkers of aging, known as epigenetic clocks, measure biological age based on DNA methylation patterns at specific CpG sites, correlating with chronological age and capturing age-related epigenome changes [35]. First-generation biomarkers, like the Horvath clocks [36,37] and Hannum clock [38], were trained using penalized regression techniques on genome-wide DNA methylation data, with Horvath clocks using multiple tissue types and cell lines, and the Hannum clock focusing on blood samples. Deviations from predicted chronological age indicate accelerated or decelerated biological aging. Second-generation biomarkers, such as GrimAge [39], PhenoAge [40], and OMICmAge [41], improved predictive power by incorporating diverse data sets and advanced algorithms, with PhenoAge including clinical measures and biomarkers, GrimAge integrating DNA methylation with surrogate biomarkers for morbidity and mortality, and OMICmAge leveraging multi-omic data. These biomarkers predict health-related phenotypes and long-term health outcomes, showing stronger associations with mortality, disease incidence, and health interventions [42].Third-generation biomarkers, such as DunedinPACE [43], use longitudinal data to measure aging pace, offering dynamic views of biological aging processes over time, and show strong associations with lifespan, healthspan, and disease onset. Vetting methods for epigenetic clocks involve assessing hazard ratios and odds ratios, correlation analyses with chronological age, mean absolute deviation calculations, and error rate assessments to ensure accurate biological age estimates across populations. Longitudinal studies or randomized controlled trials examine response to interventions by tracking changes in biological age before and after lifestyle modifications or treatments, validating the reliability, accuracy, and clinical relevance of epigenetic clocks. These biomarkers provide insights into the effects of interventions, including supplementation [44]. Research on the impact of nutritional and pharmacological supplements on epigenetic aging has focused on compounds like vitamin D and omega-3 fatty acids. Vitamin D supplementation is linked to modifications in epigenetic regulation of inflammation and metabolism genes, potentially slowing biological aging [45]. Omega-3 fatty acids, known for anti-inflammatory properties, show potential in reducing age-related methylation changes at loci associated with chronic diseases [46]. These findings suggest targeted supplementation could beneficially influence epigenetic biomarkers of aging, extending healthspan and mitigating age-associated decline.

While the individual ingredients of the Cel system have been shown to promote healthy aging phenotypes, it is not well understood whether the overall Cel system positively impacts human health, as measured by clinical lab testing and epigenetic age measures. Therefore, in this study, samples from 51 individuals over a 1 year period were used, collecting data at baseline, 3 months, 6 months, and 12 months. In addition to evaluating the Cel System supplement range, our study protocol encouraged participants to engage in 10 minutes of walking and 5 minutes of mindfulness daily. This inclusion aimed to provide a holistic approach to health improvement, combining the biochemical benefits of the supplements with the physical and mental advantages of light exercise and mindfulness practices. By integrating these activities, this study can confidently attribute the observed results to a synergistic effect of the Cel System range and the lifestyle modifications. Our primary aim was to assess the impact of the Cel System nutraceutical range on epigenetic methylation patterns, inflammation, physical performance, body composition, and epigenetic biomarkers of aging. Additionally, this study investigated how the Cel System nutraceutical range influences immune cell subsets and other epigenetic biomarker proxies (EBPs).

## RESULTS

### Participant Demographics

This study explored the potential effects of Cel System supplement range on both the clinical health and epigenetic age acceleration of healthy patients under continuous supplementation for 12-months. The analyzed cohort consisted of 51 participants (26 males and 25 females) with chronological ages ranging from 54.34 to 84.00. The study design included the longitudinal analysis of DNA methylation levels from blood and sputum samples taken at baseline, 3 months, 6 months, and 12 months to calculate epigenetic age. In tandem, clinical assessments were conducted to evaluate changes in inflammation, physical performance, and body composition markers across the same four distinct time points. The demographic information of the patients involved in the study are detailed in Table 1.

**Table 1:**
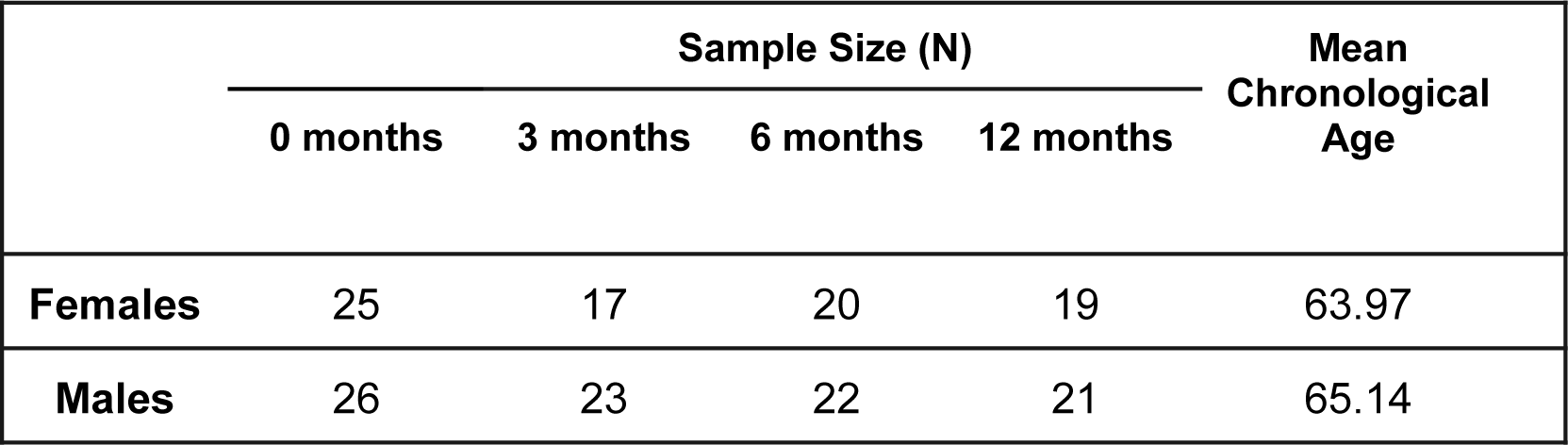
General Demographic Information of Participants.

Baseline ages of the study participants were observed to those in TruDiagnostic and Mass General Brigham’s Aging Biobank Cohorts (MGB-ABC) described in previous publications. . We compared the Epigenetic Age Acceleration (EAA) metrics calculated using three top epigenetic clocks: OMICmAge, PCGrimAge, and SystemsAge. The analysis was conducted by merging both datasets to ensure a fair comparison, and the individuals from the MGB-ABC cohort were subset to the same chronological age range (54-84) as those in the SRW cohort. For OMICmAge acceleration, the SRW cohort exhibited a mean EAA of -2.30, while the MGB-ABC cohort had a mean EAA of +0.20. The t-test yielded a highly significant p-value of 4.1x10^-9, indicating a substantial difference between the two groups. SystemsAge acceleration showed a dramatic contrast, with the SRW cohort having a mean EAA of -5.71 and the MGB-ABC cohort at +0.49. The t-test for this metric produced an extremely significant p-value of less than 2.2x10^-16. In the case of PCGrimAge acceleration, the mean EAA for the SRW cohort was -0.29, compared to +0.02 for the MGB-ABC cohort. Although this difference appeared smaller, the p-value of 0.2121 suggests that it was not statistically significant. These findings clearly demonstrate that the SRW cohort exhibits lower EAA values across multiple metrics compared to the general population, underscoring its overall healthier status.

### Cel System supplement range improves muscle strength and body function

This study assessed short term and long term physical performance outcomes associated with the Cel System supplement range. A significant increase following supplement intake in both grip strength (p= 0.0038, Figure 1A) and chair stand test scores (p= 1.7x10^-7^, Figure 1B) from baseline to 12 months was observed. Specifically, chair stand test scores improved at every time point, suggesting that the supplement may have an unvarying positive effect on lower body strength. By contrast, Touch Toes test performance worsened from the start till the end of the study (p = 0.036, Figure 1C). These results may be suggestive of the positive impact of long term supplementation of the Cel System supplement range on muscle strength and lower body function while also highlighting potential its negative effects on body flexibility and mobility.

**Figure 1:**
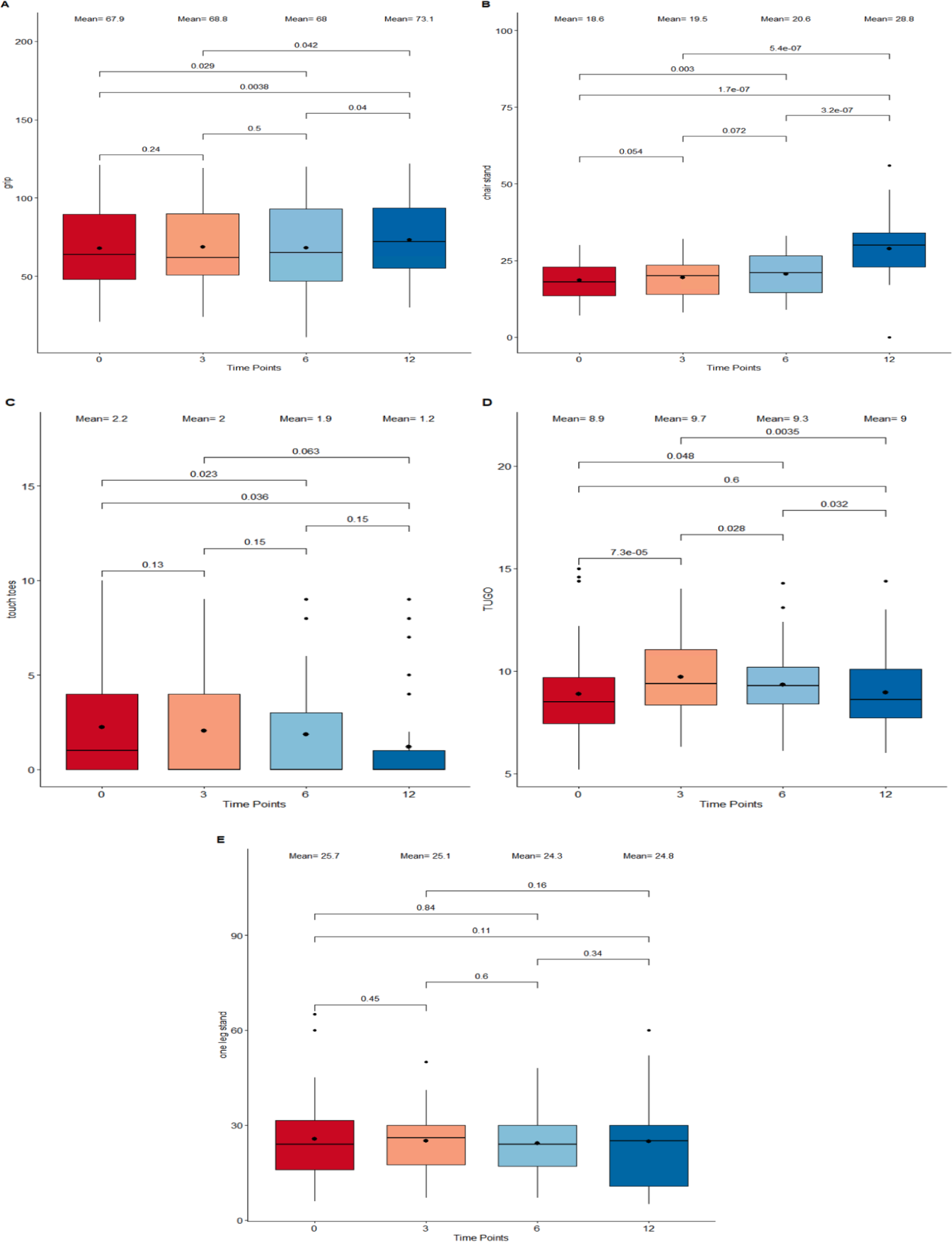
Boxplots displaying different performance marker scores of participants following supplementation. (A) Grip Strength Test. (B) Chair-Stand Test. (C) Touch-Toes Test. (D) Time Up and Go Test (TUGO). (E) One Leg Stand Test. The x-axis refers to the four time points (0 months, 3 months, 6 months and 12 months) at which the measurements were taken. The boxes represent the physical performance score outcomes which fall into the 25th to 75th percentile while all outliers are plotted as individual dots. Both mean and median at each time point are displayed as a bold dot and a horizontal line inside the box. A Wilcoxon signed-rank test was used to compare scores between adjacent and non-adjacent time points. Any change with p-value < 0.05 was considered significant.

When comparing performance marker scores between adjacent time points (baseline to 3 months, 3 months to 6 months, and 6 months to 12 months), no abrupt changes were detected. An initial mean Time Up and Go test scores of 8.9 was recorded, which peaked at 3 months (mean = 9.7; Figure 1D) and then gradually dropped between time points 3 and 6 months (p = 0.028) as well as between 6 and 12 months (p = 0.032) to reach a final mean score of 9. This implies an increment during the first 3 months that is reduced until getting the same value as the beginning after 12 months. Throughout the study duration, the one leg stand test scores remained consistent (Figure 1E), showing no significant fluctuations or variations. This observed outcome insinuates that the supplement may not affect balance.

### Cel System supplement range has no effect on inflammation

To investigate the impact of the Cel System supplement range on inflammation, variation trends in clinical measures of C-Reactive Protein (CRP) and Interleukin-6 (IL-6) levels were analyzed. The paired comparisons between the start and end of study as well as all the adjacent intermediate time points were considered. Results reveal no drastic change in either CRP and IL-6 levels from baseline to 12 months (Figure 2). While clinical CRP levels significantly increased from baseline to 3 months post supplementation (p = 0.019; Figure 2A), this was followed by fluctuations in measures across the time points resulting in an overall non-significant decrease from 3 months till the end of the study at 12 months. Clinical IL-6 levels remained relatively constant throughout the duration of the study with little variations between the four different time points (Figure 2B). These findings suggest that the Cel System supplement range does not have a notable effect on systemic inflammation mediated by these two markers.

**Figure 2:**
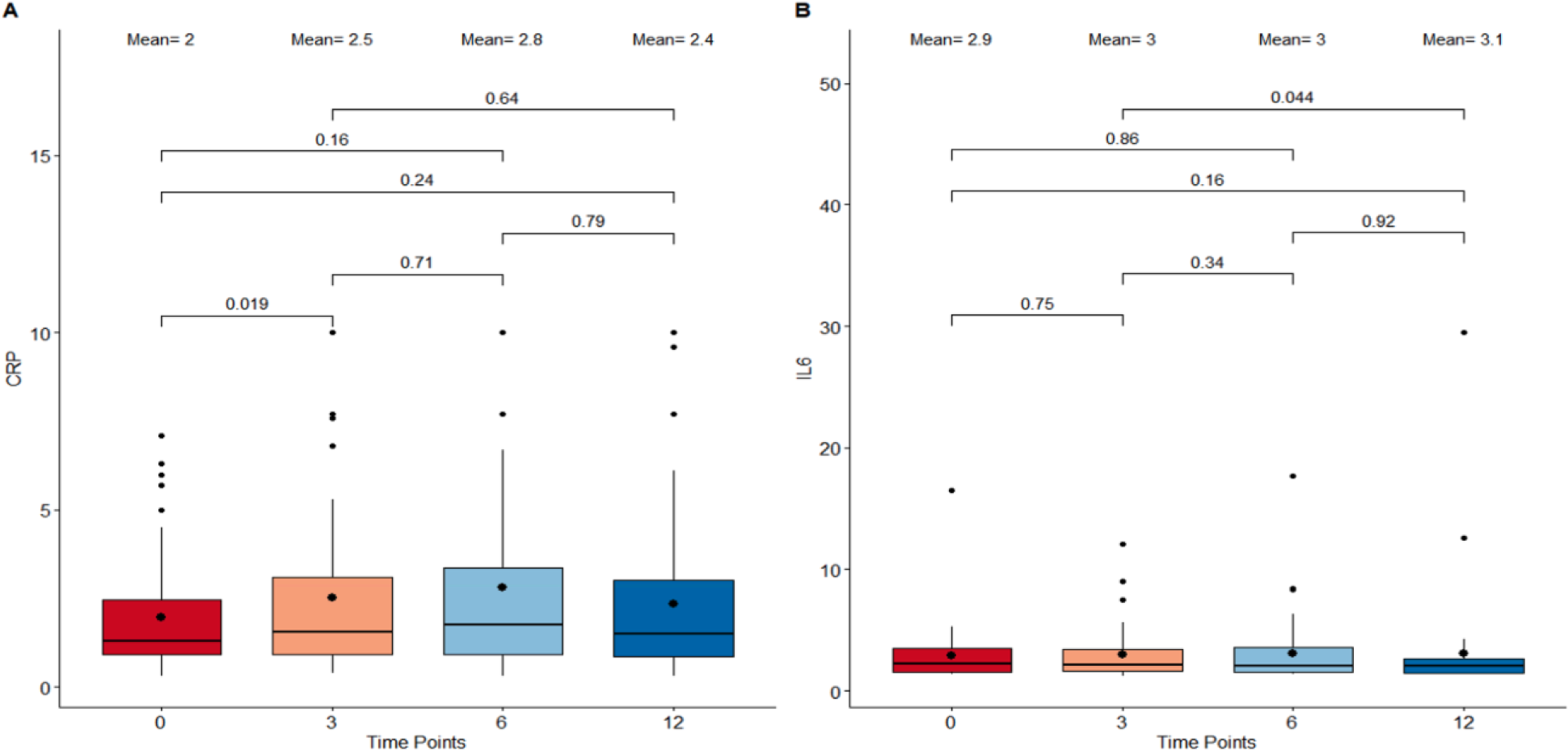
Boxplots detailing the levels of CRP and IL-6 immunological metrics following supplementation. (A) C-reactive protein (CRP) levels. (B) Interleukin-6 (IL-6) levels. The X-axis refers to the four different timepoints starting with 0 months as baseline, 3 months, 6 months, and 12 months as the final measure. The central boxes each represent the interquartile range (IQR) where 50% of the data at that time point lie. The vertical whiskers extend to the minimum and maximum values within 1.5 times the IQR from the first and third quartiles respectively. All the individual data points outside this range are considered outliers. The bold dot inside the box depicts the mean while the horizontal line indicates the median. Paired comparisons using Wilcoxon Signed-Rank Test were performed between adjacent as well as non adjacent time points and any p-value < 0.05 was considered statistically significant.

### The Cel System supplement range contributes to favorable changes in body composition metrics

Body composition is an essential component in determining the physical health and wellness of individuals. In this study, the weight, waist circumference, and body mass index (BMI) were used as metrics to explore the influence of Cel supplementation on body composition (Figure 3). Overall, notable weight loss, BMI decrease, and waist circumference reduction were found when comparing measures from baseline to the ones after 12 months of supplement intake (p = 0.033, p = 0.038, and p = 0.0018, respectively). There were no drastic variations in any measure observed during the initial 6 months of the study period. However, a significant shrinkage in the waist circumference size between the 6 month time point (mean = 38.7 in) and the end of the study (mean = 36.8 in) was observed.

**Figure 3:**
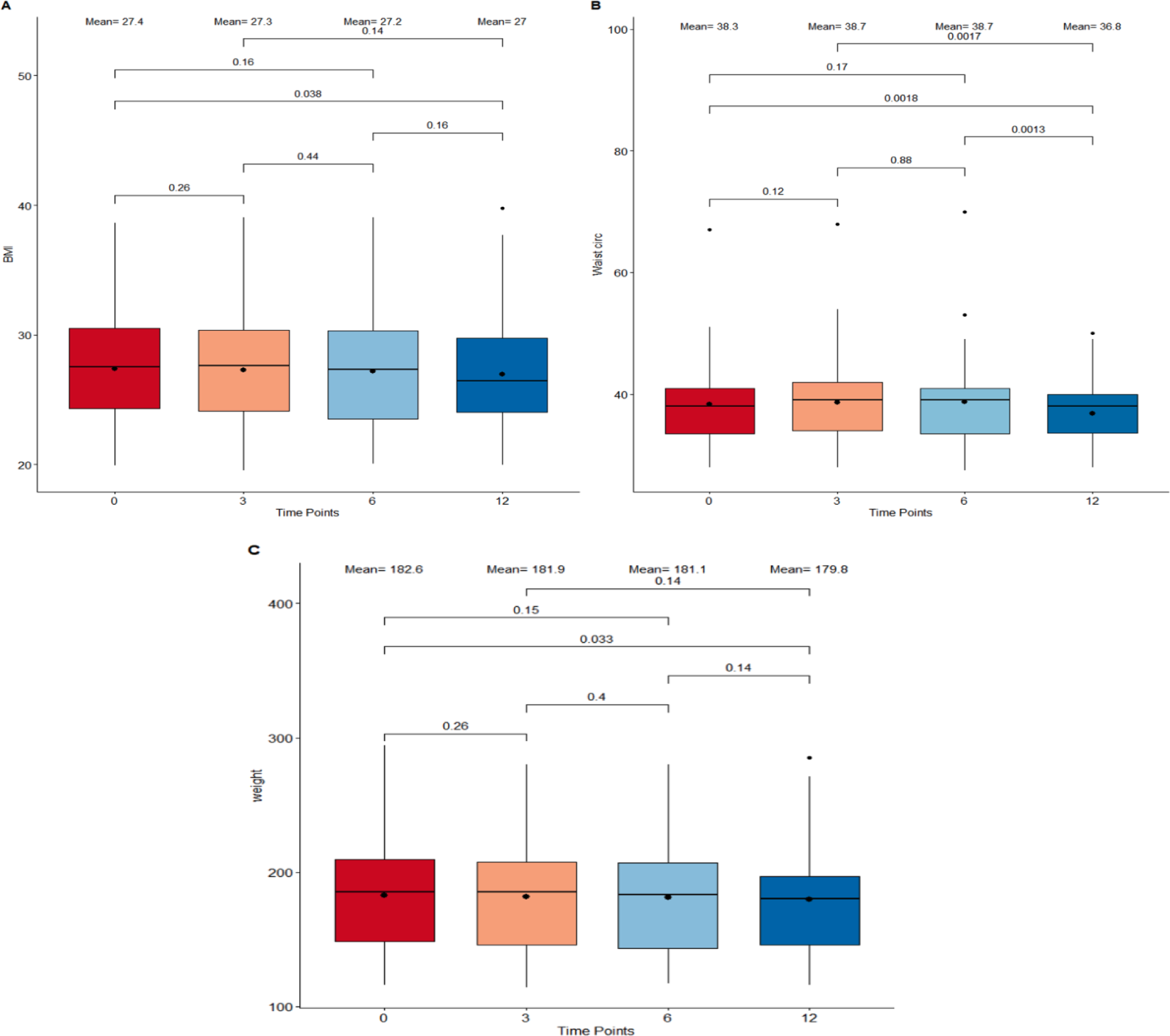
Boxplots detailing body composition measurements between timepoints. (A) Body Index Mass (BMI) in kg/m^2^. (B) Waist circumference in inches. (C) Weight in pounds. The four different time points : 0 months, 3 months, 6 months, and 12 months are plotted on the X-axis. The boxes correspond to all measurements which fall within the 25 to 75% interquartile range. The means at each time point as well as the median are displayed as a point and horizontal line respectively inside the box. All individual points which extend beyond the upper and lower limits of the IQR box plot and vertical whiskers are outlier measurements. All paired comparisons between adjacent and non-adjacent timepoints with a p-value <0.5 were considered statistically significant.

### Cel System supplement range influences epigenetic age

In order to obtain better insight on how the Cel System supplement range affects age associated with molecular changes, the biological age of participants was calculated using multiple epigenetic biomarkers of aging at baseline, 3 months, 6 months, and 12 months. The quantification of epigenetic age predictors was achieved through first generation clocks (the principle component -PC- based multi-tissue Horvath, skin and blood Horvath, Hannum, Stochastic clocks and Intrinclock), multiple second generation clocks (PhenoAge, GrimAge, Retroclock, OMICmAge, Marioni cAge, and DNAmTL), the third generation clock DunedinPACE, fitness epigenetic biomarkers, and three causal biomarkers of aging (CausAge, DamAge, and AdaptAge). To compare the metrics in different timepoints, the epigenetic age acceleration (EAA) was calculated by regressing out chronological age and technical batches.

After 12 months of the Cel System supplement range consumption, significant changes in multiple biomarkers of aging (Table 2) were observed. Notably, the epigenetic ages calculated by the first generation clock PC Horvath pan-tissue and the DNAmGrip biomarker decreased significantly at 12 months compared to baseline (p = 0.048, and p = 0.019, respectively). In contrast, PC Horvath skin and blood and third generation clock DunedinPACE exhibited a increase at 12 months compared to baseline (p = 0.045 and p = 7.4 x10^-5^, respectively). While other various age metrics showed no overall significant changes between the start and end of the study, intermediate fluctuations in clock measures were found. Remarkably, OMICmAge, Retroclock (version 2) and DNAm Fitness Age all recorded significant accelerated aging at 3 months relative to baseline (p = 0.013, p = 0.011, and p = 0.013, respectively). In line with the increment at 3 months on DNAm Fitness Age, DNAm FEV1 was reduced during this period of time (p = 0.0017).Notably, significant reductions at 3 months and 6 months were observed compared to baseline in the DamAge measurements known to assess the aging process in relation cellular damage accumulation (p = 0.0029 and p = 0.0014, respectively). Epigenetic age measured by PC Hannum, PC GrimAge, CausAge, IntrinClock, and Stochastic PhenoAge initially increased and peaked at 6 months, after which it significantly dropped in the final 6 months. Other age metrics, including PC PhenoAge, DNAm telomere length, AdaptAge, version 1 of Retroclock, DNAm Gait and VO2max, marioni cAge, and Zhang and Horvath Stochastic biomarkers, remained unchanged following supplementation.

**Table 2.** Statistical comparisons of epigenetic age acceleration (EAA) between baseline, 3 months, 6 months, and 12 months of different aging biomarkers. The first column reports the different epigenetic biomarkers of aging assessed. The next 4 columns refer to the EAA means of each clock at each time point (0 months, 3 months, 6 months, and 12 months). The final 5 columns are the p-values obtained from a Wilcoxon-Signed Rank Test comparison between adjacent and non adjacent timepoints.

|  | Mean |  |  |  | P-value |  |  |  |  |  |
| --- | --- | --- | --- | --- | --- | --- | --- | --- | --- | --- |
|  | Baseline | 3 months | 6 months | 12 months | Baseline vs 3 months | 3 months vs 6 months | 3 months vs 12 months | Baseline vs 6 months | 6 months vs 12 months | Baseline vs 12 months |
| **PCHorvath pan tissue** | 0.6 | **-0.09** | **-0.36** | **-0.15** | **0.041\*** | 0.5 | 0.85 | **0.00061\*** | 0.41 | **0.048\*** |
| **PCHorvath skin and blood** | -1.23 | **0.44** | **0.52** | **-0.31** | **9.9x10<sup>-5</sup>\*** | 0.6 | 0.11 | **7.7x10<sup>-5</sup>\*** | **0.018\*** | **0.045\*** |
| **PCHannum** | -0.45 | 0.27 | **0.31** | -0.15 | 0.052 | 0.61 | 0.66 | **0.027\*** | 0.28 | 0.8 |
| **PCPhenoAge** | -0.07 | -0.08 | 0.24 | -0.21 | 0.75 | 0.48 | 0.92 | 0.44 | 0.26 | 0.96 |
| DunedinePACE | 0.94 | 0.95 | 0.95 | **0.99** | 0.15 | 0.38 | **0.0054\*** | 0.28 | **0.0036\*** | **7.4x10<sup>-5</sup>\*** |
| OMICmAge | -0.34 | **0.58** | 0.31 | 0.17 | **0.013\*** | 0.43 | 0.31 | 0.071 | 1 | 0.13 |
| PCGrimAge | 0.02 | 0.35 | 0.37 | **-0.02** | 0.11 | 0.82 | 0.062 | 0.068 | **0.017\*** | 0.62 |
| PCDNAmTL | -9.0x10<sup>-4</sup> | 7x10<sup>-4</sup> | 0.0043 | 0.006 | 0.96 | 0.62 | 0.79 | 0.73 | 0.72 | 0.3 |
| CausAge | -0.48 | -0.38 | **0.55** | -0.17 | 0.69 | 0.12 | 0.96 | **0.039\*** | 0.22 | 0.61 |
| DamAge | 2.46 | **-2.07** | **-1.13** | -0.12 | **0.0029\*** | 0.6 | 0.19 | **0.0014\*** | 0.43 | 0.12 |
| AdaptAge | -1.12 | 0.64 | 0.26 | -0.2 | 0.17 | 0.95 | 0.75 | 0.41 | 0.69 | 0.54 |
| Retroclock v1 | -0.43 | -0.25 | -0.24 | -0.04 | 0.56 | 0.85 | 0.95 | 0.73 | 0.86 | 0.68 |
| Retroclock v2 | -0.61 | **0.41** | **0.52** | -0.02 | **0.011\*** | 0.55 | 0.3 | **0.0022\*** | 0.11 | 0.4 |
| Intrinclock | 0.151 | 0.57 | 0.448 | **-1.247** | 0.35 | 0.65 | **0.017\*** | 0.64 | 0.063 | 0.065 |
| DNAmFitAge | -0.92 | **0.45** | -0.23 | -0.35 | **0.013\*** | 0.42 | 0.18 | 0.24 | 0.99 | 0.44 |
| DNAmGait | 0.01 | -0.005 | 0.002 | 0.01 | 0.13 | 0.55 | 0.18 | 0.35 | 0.13 | 0.55 |
| DNAmGrip | 1.43 | 1.25 | 1.23 | **0.68** | 0.51 | 0.48 | 0.06 | 0.56 | 0.14 | **0.019\*** |
| DNAmVO2max | 0.34 | -0.14 | 0.32 | 0.08 | 0.098 | 0.24 | 0.37 | 0.77 | 0.72 | 0.41 |
| DNAmFEV1 | 0.04 | **0.01** | 0.02 | 0.02 | **0.0017\*** | 0.24 | 0.3 | 0.071 | 0.92 | 0.47 |
| Marioni cAge | -0.2 | 0.1 | 0.36 | -0.02 | 0.5 | 0.44 | 0.75 | 0.19 | 0.34 | 0.64 |
| Stochastic Zhang | 0.34 | 0.28 | 0.08 | 0.34 | 0.77 | 0.88 | 0.54 | 0.4 | 0.48 | 0.79 |
| Stochastic Horvath | -0.04 | 0.69 | 0.02 | 0.52 | 0.31 | 0.28 | 0.9 | 0.93 | 0.48 | 0.72 |
| Stochastic PhenoAge | -1.02 | **0.34** | **2.09** | **0.02** | **0.034\*** | **0.0128\*** | 0.77 | **0.00015\*** | **0.034\*** | 0.44 |

Using the SystemsAge approach, the independent biological age changes of different organ systems following Cel System supplement range intake were investigated. Among all the systems, blood, brain, inflammation, hormone, immune, liver, metabolic, musculoskeletal, and the overall SystemsAge had a similar trend, where the maximum level was identified at 3 months and it was followed by a deceleration between 3 and 12 months (Table 3). Kidney and heart systems had a similar trend but the highest level was at 6 months. Oppositely, the lung system showed a reduction between baseline and 12 months (p = 0.0061) with no significant increments in the intermediate timepoints.

**Table 3.** Statistical epigenetic age acceleration (EAA) comparisons of different organ systems between baseline, 3 months, 6 months, and 12 months. The different systems assessed by the SystemsAge clock are outlined in the first column along with the marginal mean values at each of the four timepoints. A paired Wilcoxon-Signed Rank test was performed between both adjacent and nonadjacent timepoints and all resulting p-values were included in the final 5 columns. P-values <0.05 were considered statistically significant.

|  | Mean |  |  |  | P-value |  |  |  |  |  |
| --- | --- | --- | --- | --- | --- | --- | --- | --- | --- | --- |
|  | Baseline | 3 months | 6 months | 12 months | Baseline vs 3 months | 3 months vs 6 months | 3 months vs 12 months | Baseline vs 6 months | 6 months vs 12 months | Baseline vs 12 months |
| **SystemsAge** | -0.72 | **0.08** | 0.07 | **-0.49** | **0.029\*** | 0.73 | 0.23 | 0.071 | **0.012\*** | 0.73 |
| **Blood** | -0.99 | **0.47** | 0.1 | -0.33 | **0.005\*** | 0.64 | 0.59 | 0.0718 | 0.3 | 0.18 |
| **Brain** | -1.12 | **0.66** | **0.66** | -0.36 | **0.0012\*** | 0.6 | 0.22 | **0.0069\*** | 0.091 | 0.3 |
| **Inflammation** | -0.46 | **0.69** | 0.08 | -0.22 | **0.014\*** | 0.3 | 0.25 | 0.3 | 0.4 | 0.57 |
| **Heart** | -0.89 | **0.13** | **0.32** | **-0.48** | **0.0065\*** | 0.57 | 0.15 | **0.011\*** | **0.0044\*** | 0.48 |
| **Hormone** | -0.42 | -0.01 | **-0.73** | -0.21 | 0.12 | **0.036\*** | 0.99 | 0.54 | 0.38 | 0.56 |
| **Immune** | -0.27 | 0.4 | 0.04 | -0.15 | 0.41 | 0.61 | 0.96 | 0.55 | 0.76 | 0.57 |
| **Kidney** | -0.59 | **0.43** | 0.46 | -0.09 | **0.041\*** | 0.76 | 0.72 | 0.06 | 0.12 | 0.59 |
| **Liver** | -1.48 | **0.73** | **0.56** | **-0.27** | **9.1x10<sup>-5</sup>\*** | 0.98 | 0.26 | **0.0013\*** | 0.06 | **0.034\*** |
| **Metabolic** | -0.91 | **0.81** | 0.11 | -0.21 | **0.0028\*** | 0.34 | 0.22 | 0.16 | 0.29 | 0.29 |
| **Lung** | 0.56 | **-0.53** | **-0.37** | **-0.45** | **0.0073\*** | 0.57 | 0.51 | **0.019\*** | 0.32 | **0.0061\*** |
| **MusculoSkeletal** | -0.96 | **0.74** | **0.6** | -0.23 | **0.00013\*** | 0.61 | 0.12 | **8.2x10<sup>-5</sup>\*** | 0.084 | 0.17 |

### The Cel System supplement range influences overall stem cell division rate

The mitotic clock epiTOC2 was utilized to look into the impact of supplementation on the mitotic rate and overall proliferative activity of stem cells. Based on the total number of stem cell replication cycles (tnsc, Figure 4A) estimated, results show a lower stem cell turnover rate at 12 months compared to baseline (p= 0.024). Specifically, the largest decrease in cumulative number of cell divisions was observed between 0 months and 3 months (p= 0.00018). Similar trends were observed for the intrinsic stem cell division cycles (irS, Figure 4B), where a significant deceleration in intrinsic proliferation rate was also prominent following 12 months of supplementation (p= 0.027) and most significant between 3 months compared to baseline (p=0.00016).

**Figure 4.**
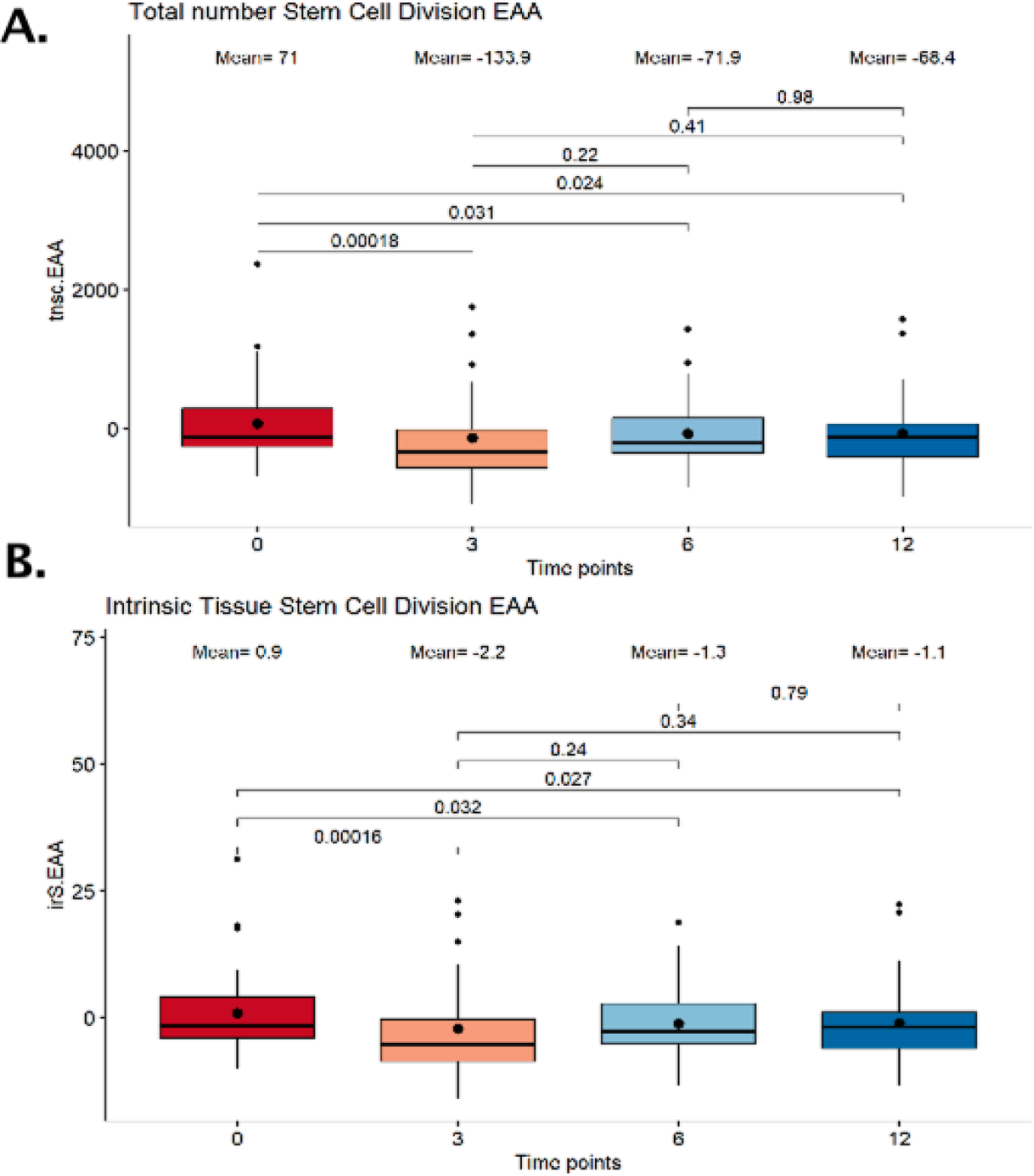
Boxplot showing the estimated number stem cell divisions following supplementation. (A) Total number of stem cell division (tnsc). (B) Intrinsic tissue stem cell divisions (irS). The x-axis depicts 4 different time points: 0 months, 3 months, 6 months, and 12 months. The boxes at each timepoint encompasses 50% of the estimated values. All outlier measures exceeding 1.5 times the interquartile range delimited by the vertical whiskers are displayed as individual points. The median and means are represented inside the boxes as a horizontal line and bold dot respectively. The stem cell division numbers between adjacent and nonadjacent time points were compared by a paired Wilcoxon Signed-Rank test and all p-values <0.05 were considered significant.

### The Cel System supplement range modulates immune cell levels

Understanding the impact of continuous supplement intake on the immune system is essential in unraveling cellular and molecular relationships between specific dietary nutrients and aging processes associated with immunity. For this purpose, variations in 12 immune cell percentages were analyzed. CD4T naive cells, CD4T memory cells, CD8T naive cells, CD8T memory, B memory, B naive, T regulatory, basophil, natural killer, neutrophil, eosinophil, and monocyte whole blood levels were all quantified at each time point using the Epidish (2023) frame (Table 4).

**Table 4.** Statistical comparisons of 12 ¡immune cells at 0 months, 3 months, 6 months and 12 months. The first four columns show the mean values of every immune cell subtype at each time point while the following columns contain the p-values of the paired-test performed between adjacent and nonadjacent time point measures. P-values <0.05 were considered statistically significant.

|  | Mean |  |  |  | P-value |  |  |  |  |  |
| --- | --- | --- | --- | --- | --- | --- | --- | --- | --- | --- |
|  | Baseline | 3 months | 6 months | 12 months | Baseline vs 3 months | 3 months vs 6 months | 3 months vs 12 months | Baseline vs 6 months | 6 months vs 12 months | Baseline vs 12 months |
| CD4T Naive | 0.004 | -0.001 | 0.001 | 0.003 | 0.18 | 0.86 | 0.62 | 0.1 | 0.77 | 0.37 |
| CD4T Memory | 0.007 | **0.006** | 0.003 | 0.002 | **0.0029\*** | 0.57 | 0.35 | 0.057 | 0.66 | 0.57 |
| CD8T Naive | 0.001 | **$3.0 \times 10^{-4}$** | **$7.0 \times 10^{-5}$** | **$2.0 \times 10^{-4}$** | **0.00012\*** | **0.017\*** | **0.02\*** | **$1.4 \times 10^{-4}$ \*** | **0.0054\*** | **$3.5 \times 10^{-5}$ \*** |
| CD8T Memory | -0.002 | -0.002 | -0.002 | -0.004 | 0.61 | 0.86 | 0.36 | 1 | 0.54 | 0.49 |
| B Naive | -0.0001 | 0.001 | 0.0003 | 0.001 | 0.22 | 0.41 | 0.89 | 0.96 | 0.92 | 0.37 |
| B Memory | 0.002 | **-0.001** | $6.0 \times 10^{-5}$ | 0.001 | **0.045\*** | 0.48 | 0.44 | 0.12 | 0.33 | 0.25 |
| T-Regulatory | 0.003 | **-0.003** | -0.001 | -0.001 | **0.021\*** | 0.64 | 0.69 | 0.12 | 0.62 | 0.19 |
| Basophils | -0.0003 | 0.0007 | $-1 \times 10^{-5}$ | $1 \times 10^{-5}$ | 0.11 | 0.45 | 0.82 | 0.8 | 0.72 | 0.6 |
| Natural Killer | 0.0003 | 0.001 | -0.0003 | 0.001 | 0.71 | 0.75 | 0.32 | 0.54 | 0.57 | 0.37 |
| Neutrophils | -0.013 | 0.005 | 0.005 | -0.003 | 0.71 | 0.8 | 0.92 | 0.25 | 0.51 | 0.75 |
| Monocytes | 0.001 | 0.003 | -0.0003 | 0.0004 | 0.4 | 0.47 | 0.44 | 0.72 | 0.8 | 0.82 |
| Eosinophils | -0.0004 | **0.001** | **0.0001** | **0.0005** | **0.00018\*** | **0.00056\*** | **0.0013\*** | **0.0069\*** | **0.00066\*** | **0.0012\*** |

Results revealed noticeable changes in several immune cell subtype proportions. While B memory, CD4 T memory and T-regulatory cells insignificantly increased in the latter time points of the study, all 3 immune cell levels initially showed a striking decline between baseline and 3 months (p = 0.045, p = 0.00012, p = 0.021, respectively). Eosinophils displayed a more gradual, yet significant increase in whole blood percentage after 12 months of supplementation (p = 0.0012). This was also the case for CD8T cells, whose levels were categorized by a proportional reduction between the start and end of the study (p = 3.5x10^-5^). Other immune subset levels did not vary between baseline and 12 months.

### The Cel System supplement range affects Epigenetic Biomarker proxies and Marioni EpiSign scores for proteomic prediction

In efforts to complement the above epigenetic age measures and provide a more comprehensive reflection of the physiological, proteomic, and molecular changes associated with the Cel System supplement range, 396 previously developed DNAm-based epigenetic biomarker proxies (EBPs) were first calculated at all intermediate time points and compared means between baseline and 12 months. Results showed significant changes in 2 EBP predicted measures, deoxycholic acid glucuronide and Transthyretin (TTHY). However, in terms of directionality, these metabolic biomarkers followed opposite trends. While deoxycholic acid glucuronide, a bile acid byproduct related to liver function decreased following 12 months of supplementation, TTHY, a retinol and thyroid hormone transport protein, increased. All EBP mean values and start to end comparisons are detailed in Supplementary Table 1.

Next, Marioni protein biomarker levels were analyzed to see how they differ between baseline and 12 months of the Cel System supplement range dietary enhancement (Supplementary Table 2). Among all the biomarkers assessed, 2 protein estimates with varying means were identified (Supplementary Table 2). The inflammatory chemokine CCL25, typically associated with intestinal epithelial cells, and the Hepatocyte Growth Factor Activator, involved in hepatic tissue regeneration and repair, both demonstrated noteworthy reductions.

### The Cel System supplement range impacts whole-genome DNA methylation levels

To assess the overall change in global DNA methylation patterns following Cel System supplementation, an epigenetic-wide association study (EWAS) was conducted and differentially methylated gene loci for all the CpG sites across the genome were identified. After adjusting for multiple confounding factors such as age, sex, cell type proportions, and other technical variables, a total of 1655 differentially methylated loci were found. Among these significant CpG sites, 674 loci were hypermethylated and 685 sites were hypomethylated when comparing baseline to 12 months (FDR <0.05). The Manhattan plot (Figure 6) reveals the unique genes these differentially methylated loci map to while Table 5 details the top 20 differentiated CpG sites.

**Figure 6.**
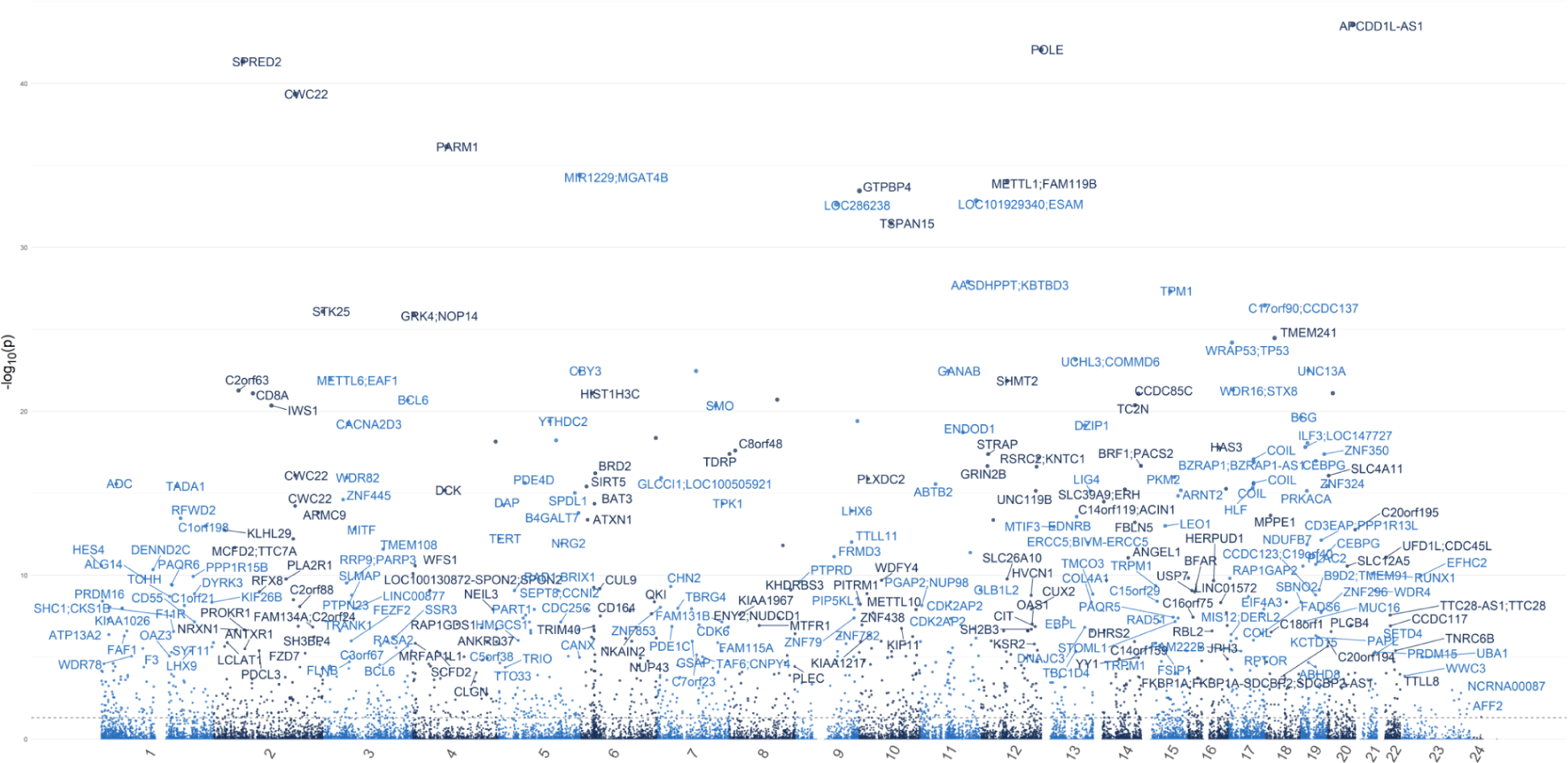
Manhattan Plot displaying the overall genomic distribution of unique genes associated with differentially methylated CpG sites. Individual dots represent different CpG sites with the x-axis denoting their color-coded chromosomal location. The y-axis indicates the negative logarithm (base 10) of the p-value. A False Discovery Rate (FDR) lower than 0.05 was considered as a significant threshold.

**Table 5.**
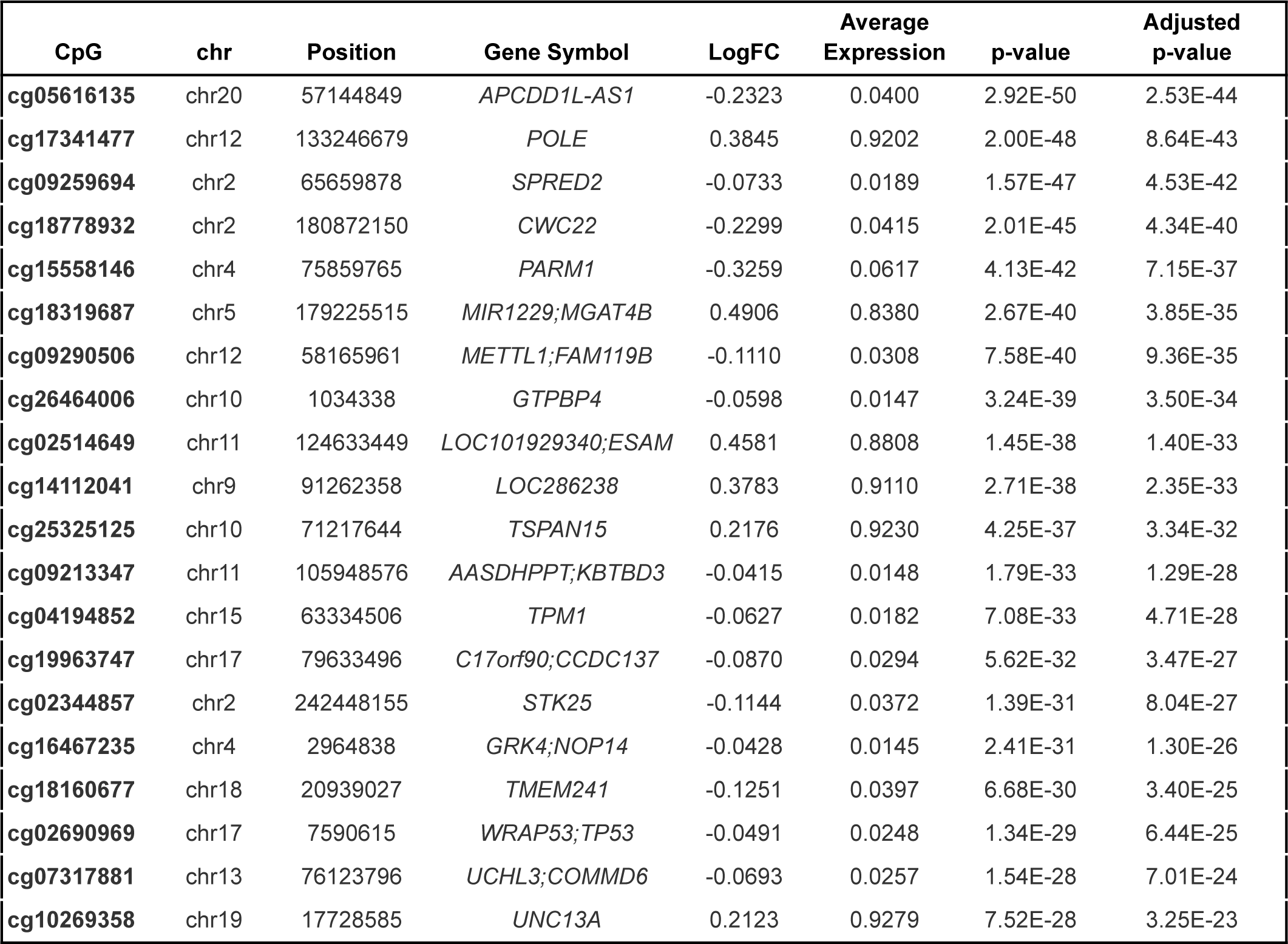
Top 20 differentially methylated CpG sites following supplementation. Each CpG site is identified by its chromosomal location, base pair position, and associated gene symbol (when it is not an intergentic CpG site). The mean log fold change (LogFC) denotes the directionality and magnitude of the methylation change across the timepoint 4 (12 months) compared to baseline, with negative values indicating hypomethylation and positive values indicating hypermethylation. This is followed by the mean expression levels of each CpG site across all four timepoints. The last two columns report both nominal p-values and the False Discovery Rate (FDR) adjusted p-values.

To further identify functional implications in relation to biological processes, molecular functions and cellular components, an enrichment analysis was performed using the differentially methylated CpG sites. Based on the direction of methylation, the GREAT analysis results show that hypermethylated sites were mostly enriched for oxidative stress-induced premature senescence, pyrimidine deoxyribonucleotide metabolic process and hyaluronan biosynthetic process (Supplementary Figure 1). As for related molecular functions, these CpG sites were highly associated with TRAIL binding hyaluronan synthase activity, and amide transmembrane transporter activity. By contrast, the enrichment of GO terms with the hypomethylated CpG loci highlighted the over-representation of neurotransmitter loading into synaptic vesicles,pore complex assembly, and collagen biosynthetic process (Supplementary Figure 2). Among the enriched molecular functions, protein phosphatase 2A binding activity and the activation of transcription factor binding were identified. While the endoplasmic reticulum lumen and collagen trimer were enriched in hypermethylated loci, the DNA-directed RNA polymerase II, holoenzyme and nuclear DNA-directed RNA polymerase complex, among others were shown to be associated with the hypomethylated CpG sites.

Methylation levels in genomic regions were compared between baseline and 12 months of supplementation. A total of 16 hypermethylated and 109 hypomethylated regions were identified with a Holm-adjusted FDR (HMFDR) < 0.05. While only 10.4% of the differentially methylated regions are intergenic, 89.6% map back to overlapping or unique genes. Notably, genes implicated in various biological processes and cellular functions such as *HAS3, SPRED2, CWC22,* and *SART1* have more than 9 significant CpG probes aligned to their locus. The top 20 genomic regions which exhibited the most methylation change between baseline and 12 months following the Cel System supplement range intake are represented in Table 6.

**Table 6.** Top 20 DNA methylation regions significant differences between 12 months following supplementation and baseline. The regions are annotated by their chromosomal location (chr), their start and end base pair position, and the number of base pairs they span (width). The number of differentially methylated CpG sites with each region as well as the genes overlapping these regions (when applicable) are also detailed. All regions with a positive mean differential coefficient are hypermethylated regions while regions with negative mean differential coefficients are hypomethylated. The last two columns list the minimum smoothed false discovery rate (FDR), and the Holm-adjusted FDR (HMFDR) which was used to determine significance.

| Chr | Start Position | End Position | Width | Number of CpG sites | Overlapping Genes | Mean Differential | Min Smoothed FDR | HMFDR |
| --- | --- | --- | --- | --- | --- | --- | --- | --- |
| chr5 | 179224563 | 179226186 | 1624 | 12 | *MGAT4B, MIR1229* | 0.0378 | 1.24E-39 | 5.21E-19 |
| chr16 | 69140794 | 69141478 | 685 | 11 | *HAS3* | -0.0063 | 1.73E-18 | 4.04E-12 |
| chr17 | 43972306 | 43973522 | 1217 | 10 | *MAPT, MAPT-AS1* | -0.0134 | 1.26E-22 | 5.53E-12 |
| chr2 | 65659148 | 65660700 | 1553 | 12 | *SPRED2* | -0.0151 | 2.12E-18 | 3.62E-11 |
| chr16 | 56965376 | 56966475 | 1100 | 12 | *HERPUD1* | -0.0041 | 2.63E-17 | 4.52E-11 |
| chr22 | 19466331 | 19467326 | 996 | 15 | *CDC45, UFD1L* | -0.0059 | 5.39E-23 | 7.43E-11 |
| chr10 | 1033582 | 1035006 | 1425 | 15 | *GTPBP4, AL359878.1* | -0.0049 | 1.28E-14 | 9.76E-11 |
| chr4 | 2964097 | 2965549 | 1453 | 15 | *GRK4, NOP14* | -0.0105 | 4.96E-19 | 1.42E-10 |
| chr11 | 105947676 | 105949106 | 1431 | 23 | *AASDHPPT, KBTBD3* | -0.0053 | 1.03E-16 | 1.47E-10 |
| chr4 | 75859212 | 75860346 | 1135 | 6 | *PARM1* | -0.0858 | 8.28E-18 | 2.22E-10 |
| chr7 | 75795555 | 75796528 | 974 | 7 |  | -0.0152 | 6.79E-16 | 1.02E-09 |
| chr3 | 15468722 | 15469386 | 665 | 12 | *EAF1, METTL6* | -0.0113 | 9.94E-18 | 1.18E-09 |
| chr16 | 11438416 | 11439785 | 1370 | 13 | *RMI2, RP11-485G7.5* | -0.0106 | 3.38E-14 | 2.33E-09 |
| chr12 | 58165256 | 58166652 | 1397 | 16 | *METTL21B, METTL1* | -0.0096 | 5.30E-15 | 3.78E-09 |
| chr2 | 180871638 | 180872164 | 527 | 14 | *CWC22* | -0.0654 | 2.33E-30 | 5.67E-09 |
| chr11 | 65728931 | 65729635 | 705 | 9 | *SART1* | -0.0053 | 4.52E-11 | 1.19E-08 |
| chr11 | 94822598 | 94823603 | 1006 | 6 | *ENDOD1* | -0.0084 | 1.29E-18 | 1.44E-08 |
| chr14 | 100070556 | 100071437 | 882 | 9 | *RP11-543C4.1* | -0.0118 | 1.26E-12 | 2.79E-08 |
| chr17 | 79633202 | 79633857 | 656 | 15 | *CCDC137, OXLD1* | -0.0068 | 1.47E-09 | 3.64E-08 |
| chr15 | 63333846 | 63335016 | 1171 | 8 | *TPM1* | -0.0080 | 3.75E-15 | 3.89E-08 |

## DISCUSSION

The integration of supplements into health management has received considerable attention due to their promising ability to influence various health metrics. The expanding body of evidence indicating that certain supplements may enhance cardiovascular health, cognitive function, and physical performance has further heightened this interest. Investigating the effects of supplements may reveal how they can enhance overall health by reducing biological aging, improving immune function, supporting optimal body composition, and addressing specific nutritional deficiencies. In this study, the Cel System supplement range showed significant effects on patients who took the supplement over a one year period. In particular, supplementation resulted in improved muscle strength, body function, and body composition metrics, epigenetic changes related to biological age, impacted stem cell division rates, modulated immune cell levels, and significantly changed EBPs and Maroni biomarker proxies.

Physiological function measures are clinically meaningful as they provide a comprehensive assessment of an individual’s physical capabilities and overall health, which are crucial for understanding the impacts of aging and evaluating interventions across studies. The improvements in grip strength and chair stand test scores observed when comparing physical performance measurements across the four time points in this study are suggestive of the Cel System’s long term benefit on muscle strength and lower body function. Numerous studies have investigated both the individual ingredients of this supplement and their combined effects, revealing their efficacy in promoting cognitive, physiological, and physical benefits. For instance, Nicotinamide mononucleotide (NMN) supplementation was previously linked to improvements in muscle function and physical performance metrics such as grip strength and endurance as it plays a crucial role in supporting mitochondrial energy production in muscles and other tissues [22,47]. Likewise, Rutin and B12, both found in Cel System, have been found to enhance muscle strength, reduce muscle damage, and prevent weakness [48,49]. Vitamin C’s involvement in collagen synthesis also hints at supporting musculoskeletal health [50].

Our results also highlight the positive influences of Cel System supplement range on body composition. Continuous supplement intake was marked by weight loss, BMI decrease and waist circumference reductions which may be an indicative of the underlying metabolic changes occurring as a result of the ingredients in the Cel supplement. Specifically, Rutin has been proven to aid in weight management and lipid metabolism [51], while NMN, as a nicotinamide adenine dinucleotide (NAD+) precursor, has been implicated in regulating cellular energy metabolic pathways and enhancing insulin sensitivity [52,53]. Astragaloside has also been studied for its potential to improve metabolic health [54,55].

While the study’s findings align with weight management, BMI regulation and muscle strength improvements associated with supplement intake, this study reports no significant enhancement in flexibility and neuromuscular activity. This nuanced outcome is noteworthy, particularly when considering that in addition to the supplement consumption, participants were encouraged to practice mindfulness for 5 minutes and walk for 10 minutes on a daily basis as part of the study’s protocol. As such, highlighting both the positive and negative elements of physiological testing could provide a compelling argument that the specificity of the observed effects are more likely attributable to the supplement range rather than the lifestyle modifications alone. The lack of improvement in flexibility and neuromuscular connectivity further implies that the specific enhancements in body composition and lower body muscle strength may be a direct effect of Cel supplementation in combination to the daily physical activity or mindfulness practices.

Due to the antioxidant, anti-inflammatory, immunomodulatory, anti-carcinogenic, and biosynthetic properties of the ingredients found in Cel supplements, it becomes of considerable interest to investigate whether such metabolic composition also promotes biological health, and longevity through positive molecular alterations. Our results demonstrate a reversal of biological aging in some but not all DNAm based measures. The epigenetic age deceleration across diverse methodologies is surprising considering that most patients enrolled in this cohort were already much biologically younger than traditional biobank cohorts such as the MGB-ABC cohort used to create OMICmAge.

The antioxidant properties of some ingredients such as astragaloside and Rutin may mitigate the oxidative stress and reduce apoptosis. This has been shown to be correlated to cells that are less prone to constant cell division and differentiation [58]. This could explain the lower stem cell turnover rates observed using the mitotic clock epiTOC2. In the context of our study where supplementation was paralleled with a decrease in B memory, CD4 T memory ,T-regulatory, and CD8T cells, it is plausible that astragaloside and rutin, along with other ingredients found in Cel system may also be acting collectively to regulate immune function, potentially reducing excessive immune activity and chronic inflammation [59,60]. Zinc, Selenium and vitamin B12 have all been proven to modulate inflammation and dampen immune responses, mirroring the alterations observed in immune cell proportions [61–63].The maintenance of cellular integrity and homeostasis highly depends on a balanced immune response. All metabolic and proteomic biomarker changes following Cel system supplementation further underscore the ability of its active ingredients to target intricate molecular and cellular processes, pivotal for maintaining metabolic homeostasis and reducing the burden of age-related oxidative damage and chronic inflammation. Reductions in the chemokine CCl25 were noticed, which may be indicative of a healthier gut environment, less prone to intestinal and epithelial inflammation [64]. Similarly, the decrease in Hepatocyte Growth Factor Activator and deoxycholic acid (DCA) may be hinting at hepatic stress mitigation. Such notable reductions reflect potential improvements in overall intestinal and liver health following supplementation, decreasing the need for injury-alleviating metabolites that mediate protective mechanisms within these organs [65].

Furthermore, the identification of a substantial number of differentially methylated sites involved in antioxidation, multiple fatty acid and protein biosynthesis and transport pathways, and other biological processes, molecular functions and cellular components mirror sizable shifts in the epigenetic profile of the participants. Although more analysis needs to be done to confirm whether such alterations point towards a more youthful epigenome, these changes in methylation could provide additional molecular evidence supporting the systemic impact of supplementation on longevity and delaying age-related biological processes.

It is important to note however that some data showed initial worsening during the first three to six months before positive results emerged. This pattern suggests a potential adaptation period where the body adjusts to the supplement regimen. Further research is needed to evaluate whether these benefits continue to accrue over time or if a cycling protocol might be more effective. This phenomenon has also been observed in senolytic interventions, which initially induce stress responses before yielding long-term benefits [66]. Understanding these dynamics could optimize the efficacy of the Cel System supplement range and provide more nuanced guidelines for their use.

This study does acknowledge several limitations in this analysis. Our sample size of 51 compels future studies with larger cohorts to validate our observations, especially because not all the individuals followed through with the study until completion. Investigating multiple ingredients simultaneously complicates the interpretation of results due to potential interactions between the ingredients. This complexity can obscure the individual effects of each ingredient, making it challenging to isolate their specific contributions to the observed outcomes. The EBPs used are also predicted measures instead of actual protein, metabolite, and clinical variables measured at the laboratory, which presents an additional potential limitation. The study’s design as a non-randomized trial without a placebo control group limits its internal validity and the strength of its causal inferences. This lack of control over placebo effects makes it challenging to distinguish the genuine effects of the intervention from physical or physiological responses triggered by participants’ beliefs about receiving treatment. Being a pilot study, the research is primarily exploratory and designed to test the feasibility of the methodology rather than to produce definitive results. The preliminary nature of this pilot study will be utilized to refine and improve the design of subsequent, larger-scale studies and provide higher potential efficacy of the Cel System supplement range.

Given the evidence that Cel enhances physical performance, optimizes body composition metrics, influences epigenetic age, affects overall stem cell division rates, and impacts EBPs and Maroni biomarker proteins, this study concludes that this supplement may effectively reduce biological age. This study confirms the capabilities of the Cel System supplement range through high significance in muscle strength and body function tests, body composition metrics, epigenetic clocks, and stem cell division rates between baseline and 12 months. Further studies should leverage this data to explore the mechanistic pathways underlying these effects. This research should aim to delineate the cellular processes by which Cel modulates these physical and physiological parameters. Detailed investigations should be conducted using advanced omics technologies, longitudinal studies, and randomized controlled trials to validate and expand upon these findings. By integrating proteomics, genomics, and metabolomics, future studies can elucidate the specific biological mechanisms and potential therapeutic applications of the Cel System supplement range. Additionally, studies should consider the dose-response relationship and long-term safety profile of the Cel System supplement range to fully establish its efficacy and potential as a novel intervention for reducing biological age and enhancing human healthspan.

## MATERIALS AND METHODS

### Study Population

This cohort consisted of 51 participants; 5 individuals had 1 sample, 2 individuals had 2 samples, 11 individuals had 3 samples, and 33 individuals had 4 samples (Table 1). 49% of participants were female. The mean chronological age of the cohort was 64.57. These individuals were recruited based on the following criteria: mean and women of any ethnicity, minimum 55 years old, female subjects must be post-menopausal, participants must be able to comply with treatment plan and laboratory tests, must be able to read, write, and speak English fluently, must have a smartphone and be able to download and use the App, must have an established primary care provider, and must be willing and able to consume study supplements throughout the duration of study period.

### DNA Methylation Assessment

Whole blood was collected at baseline, 3 months, 6 months, and 12 months for DNA methylation preparation and analysis. Blood collected by the clinics was sent to TruDiagnostic labs in Lexington, KY, for DNA extraction and methylation processing. Using the EZ DNA Methylation kit (Zymo Research), 500 ng of DNA was bisulfite-converted following the manufacturer’s instructions. Bisulfite-converted DNA samples were randomly assigned to wells on the Infinium HumanMethylationEPIC BeadChip, and the subsequent steps included amplification, hybridization, staining, washing, and imaging with the Illumina iScan SQ instrument to acquire raw image intensities. Longitudinal DNA samples for each participant were assessed on the same array to mitigate batch effects. Raw image intensities were saved as IDATs for further processing.

### Collection of Clinical Inflammation Measures

Blood samples were collected simultaneously with the TruDiagnostic tests. Instead of using finger pricks, blood was drawn from the antecubital vein using a butterfly needle. A portion of the blood was then applied directly onto TruDiagnostic pads. Samples for IL-6 analysis were stored at -80°F until a batch of 15 samples was collected. These frozen samples were then sent to Quest Labs for analysis. Blood samples for high-sensitivity C-reactive protein (hsCRP) were sent to Ulta Labs/Quest on the same day they were drawn for immediate analysis.

### Physical and Body Composition Assessment

Patients’ physical and body composition measurements were conducted consistently using standardized equipment and procedures to ensure accuracy and reliability. All measurements were performed by two trained medical assistants, ensuring consistency in the assessment process.

#### Weight and Body Composition

Patients were weighed using the InBody 970 scale, a multifrequency bioelectrical impedance analyzer known for its precision in body composition analysis. The InBody 970 provides detailed metrics including weight, body fat percentage, muscle mass, and other relevant parameters.

#### Waist Circumference

Waist circumference was measured using a cloth tape measure placed at the top of the iliac crest. This anatomical landmark ensures that measurements are taken consistently at the same location across all patients.

#### Grip Strength

Grip strength was assessed using a standardized dynamometer. The same two medical assistants conducted all grip strength tests to maintain consistency in technique and results.

#### Timed Up and Go (TUG) Test

The Timed Up and Go (TUG) test was administered to evaluate patients’ functional mobility. This test measures the time taken for a patient to rise from a chair, walk three meters, turn around, walk back, and sit down.

### Statistical analyses and reproducibility

#### Deriving Estimates of Epigenetic Clocks and Methylation-Based Metrics

DNA methylation (DNAm) data was utilized to compute a range of measures collectively referred to as epigenetic clocks or aging biomarkers. These included three clocks designed to predict the chronological age of the donor: Horvath Pan Tissue [36], Horvath Skin and Blood [37], and Hannum [38]. Additionally, two clocks were employed aimed at predicting mortality: PhenoAge [40] and GrimAge [39]. Retroclock , OMICmAge [41], and a clock for measuring telomere length, DNAmTL [67] were also utilized. Furthermore, this study incorporated elastic net-based and BULP-based Zhang clocks (Zhang-EN and Zhang-BULP) and three causal clocks (CausAge, DamAge, and AdaptAge). To assess the rate of physiological integrity decline, the DundedinPACE clock was used [43]; and to measure chronological age independent of immune cells, the IntrinClock was used [68].

To compute the principal component-based epigenetic clocks for the Horvath multi-tissue clock, Horvath Skin and Blood clock, Hannum clock, PhenoAge clock, GrimAge clock, and telomere length, a custom R script was employed available on GitHub (https://github.com/MorganLevineLab/PC-Clocks). Non-principal component-based (non-PC) epigenetic metrics for Horvath, Hannum, and DNAmPhenoAge were calculated using the *methyAge* function in the *ENmix* R package. Retroclock, OMICmAge, Zhang-EN, Zhang-BULP, CausAge, DamAge, and AdaptAge were computed. The DunedinPACE clock, assessing the pace of aging, was calculated using the *PACEProjector* function from the *DunedinPACE* package on GitHub (https://github.com/danbelsky/DunedinPACE). The mitotic clocks were determined using the *epiTOC2* function from the *meffonym* package. The IntrinClock calculation was performed as previously described in literature.

Other non-epigenetic age metrics included the relative proportions of 12 immune cell subsets estimated via EpiDISH [69], 116 predictions of biochemical and lifestyle risk factors based on methylation using *MethylDetectR* [70], and 396 proxies of epigenetic biomarkers [28].

### Statistical test for comparing metrics across time points

To start the statistical comparison between each of the metrics in each time point, they were first adjusted for potential confounding factors and technical variability. The epigenetic age acceleration (EAA) metrics were calculated by regressing out the chronological and the array type.

Statistical analyses for each EAA metric were performed using paired Wilcoxon-rank sum tests across individual timepoints. Both adjacent and non-adjacent time-point comparisons were taken into account. Statistical significance was set at a p-value below 0.05.

### Epigenome-wide association study

The epigenome-wide association study (EWAS) was performed using the *limma* Bioconductor package. A differential mean analysis was performed between baseline and one-year post treatment to see whether the supplementation was associated with changes at specific loci. Based on the available covariates, the regression models were adjusted by sex, age, cell type proportions, and other technical variables. This study also set as random effect the participant ID. The model

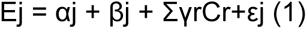

Was fitted where Ej denotes the methylation level vector across individuals at probe j (j = 1, . . . 866836), βj is the effect, Cr is the r adjusting covariate and its effect γr, and εj is the noise that follows the distribution of methylation levels with mean 0.

Adjusted P-values were calculated using FDR correction for considering multiple comparisons. The inflation or deflation of P-values across the methylome was assessed with Q-Q plots and lambda values. Significant probes were selected as those with FDR lower than 0.05 after correcting for multiple comparisons. GREAT was used to understand the functional relevance of the differentially methylated loci (DML). The GREAT software will compare genomic features against the genes of interest in order to run Gene Ontology (GO) analysis. This software looks at the number of DMLs which overlap to the promoter and enhancer regions to run a binomial enrichment analysis of identifying overrepresented/enriched GO terms.

### Differentially methylation region analysis

To evaluate the global changes on DNA methylation in regions instead of specific loci, the *DMRcate* package was used. The model was adjusted by the same covariates as the EWAS analysis: age, sex, cell type proportions, and other technical variables. Again, the DNA methylations were compared at baseline and at 12 months after the supplementation. Significant regions were considered by an adjusted p-value below 0.05.

## Author contributions

RG and VBD analyzed the data; RG, NCG, SPT, and VBD drafted the original manuscript; GM and SW recruited patients, extracted blood, and collected clinical measures; TLM generated methylation data; GM, SW, and RS designed the patient and clinical recruitment of the study; NCG and VBD designed the statistical analysis of the data; all authors proofed and edited the manuscript prior to submission.

## Conflicts of interest

VBD, RD, NCG, SPT, TLM, and RS are employees of TruDiagnostic. GM is an employee of SRW.

## Ethical Statement

This research was conducted in accordance with the ethical standards of the relevant institutional and/or national research committee(s) and with the Helsinki Declaration of 1975, as revised in 2013. All procedures involving human participants or animal subjects were approved by the appropriate ethics committee(s) and informed consent was obtained from all individual participants included in the study. The authors declare that they have no conflicts of interest that could have influenced the outcomes of this research. Any funding sources are disclosed, and the data supporting the findings of this study are available upon request.

## Funding

SRW has provided funding for data analysis and IRB funding. Regenerative Wellness provided all other costs associated with testing and patient recruitment.

## Ethical Statement

The study involving human participants was reviewed and approved by the Institute for Regenerative and Cellular Medicine (approval number IRCM-2022-336).

## Consent

The participants provided informed consent to participate in this study.

## Supplementary Materials

**Supplementary Figure 1. Gene Ontology (GO) terms enriched for the hypermethylated CpG sites after supplementation.** The Top 5 most enriched GO terms for Biological Processes (A), and Molecular Function (B) were included. A nominal p-value <0.001 was used to assign significance.

**Supplementary Figure 2. Gene Ontology (GO) terms enriched for the hypomethylated CpG sites after supplementation.** The Top 5 most enriched GO terms for Biological Processes (A), Molecular Function (B), and cellular components (C) were included.A nominal p-value <0.001 was used to assign significance.

**Supplementary Table 1. Epigenetic Biomarker Proxy (EBP) Analysis Between Baseline (0 months) and 12 Months Following Supplementation.** The first column shows the epigenetic biomarker proxy assessed. Columns 2 and 3 report the marginal mean values for each time point. The p-value column depicts the p-value of the Wilcoxon-rank sum test performed between baseline and 12 months. The final column depicts the adjusted p-value by using the false discovery rate (FDR) adjustment. The table includes 396 EBPs.

**Supplementary Table 2. Marioni protein biomarker analysis between baseline and 12 months.** The first column details all Marioni proteins assessed between baseline and 12 months. The following columns (columns 2 and 3) report the predicted marginal mean values at each of the time points. The p-value column depicts the p-value of the Wilcoxon-rank sum test performed between baseline and 12 months. The final column depicts the adjusted p-value by using the false discovery rate (FDR) adjustment.

## Notes

### Competing Interest Statement

VBD, RD, NCG, SPT, TLM, and RS are employees of TruDiagnostic. GM is an employee of SRW Laboratories.

### Summary of Updates

Author list was updated to include missing author.

