## Supplementary Figures and Tables for "The Impact of a Natural Ingredients based Intervention Targeting the Nine Hallmarks of Aging on DNA methylation"

### The effect of 12-month CEL SRW supplementation on body function and epigenetic age

**Supplementary Material**

*Supplementary Figures*

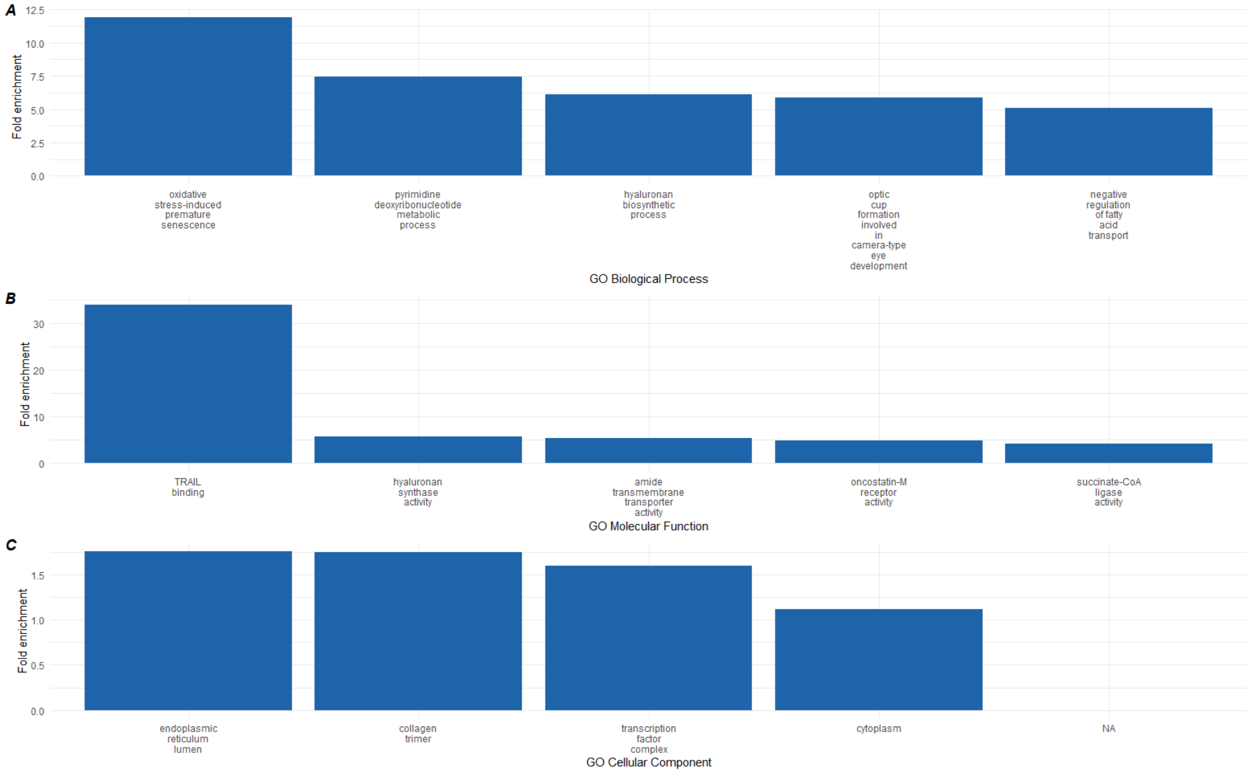


**Supplementary Figure 1. Gene Ontology (GO) terms enriched for the hypermethylated CpG sites after supplementation.** The Top 5 most enriched GO terms for Biological Processes (A), Molecular Function (B), and Cellular Component (C) were included. A nominal p-value <0.001 was used to assign significance.

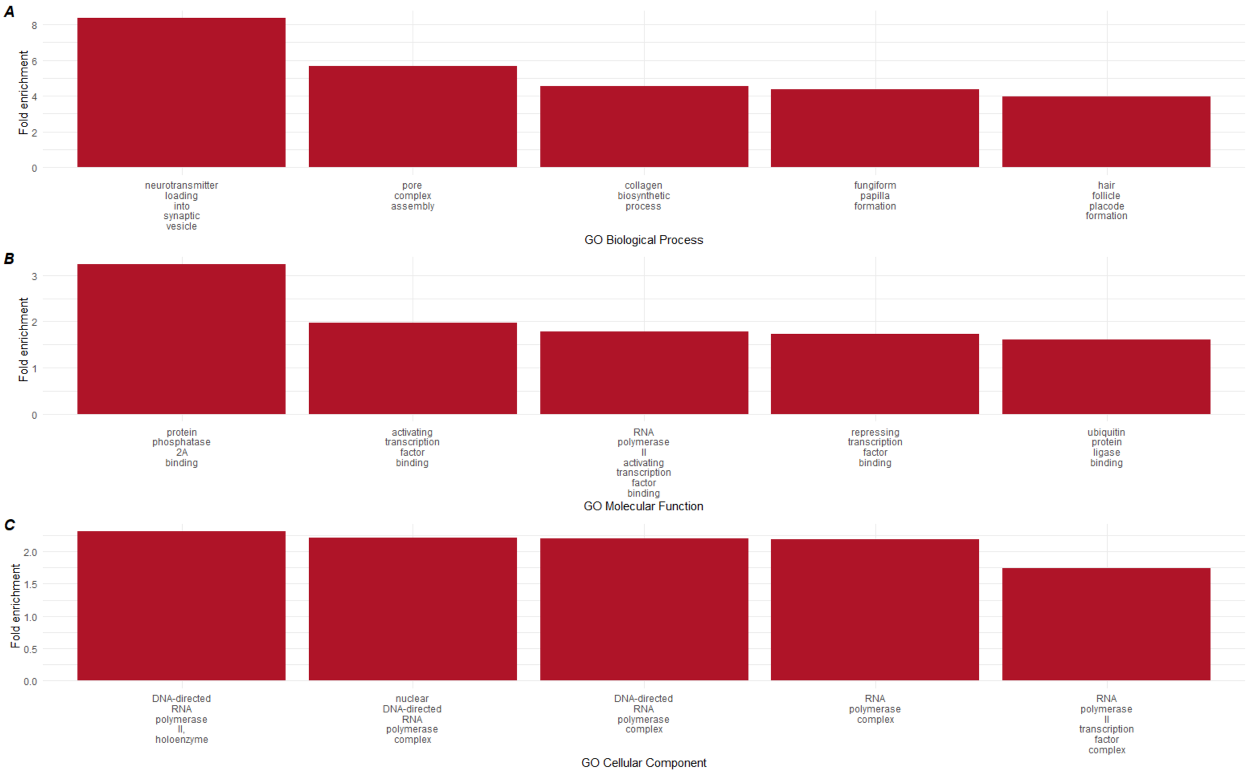


**Supplementary Figure 2. Gene Ontology (GO) terms enriched for the hypomethylated CpG sites after supplementation.** The Top 5 most enriched GO terms for Biological Processes (A), Molecular Function (B), and cellular components (C) were included.A nominal p-value <0.001 was used to assign significance.

**Supplementary Table 1. Epigenetic Biomarker Proxy (EBP) Analysis Between Baseline (0 months) and 12 Months Following Supplementation.** The first column shows the epigenetic biomarker proxy assessed. Columns 2 and 3 report the marginal mean values for each time point. These values were adjusted for confounding factors technical variability such as age, and array type. The p-value column depicts the p-value of the Wilcoxon-rank sum test performed between baseline and 12 months. The final column depicts the adjusted p-value by using the false discovery rate (FDR) adjustment. The table includes 396 EBPs.

| | Mean | | 0 vs 12 months | |
|---|---|---|---|---|
| | 0 months | 12 months | p-value | FDR |
| deoxycholic acid glucuronide | 0.09521349614 | 0.01926854981 | 0.04136160471 | 1 |
| TTHY_HUMAN | -0.01787292975 | 0.002262820717 | 0.04136160471 | 1 |
| beta-cryptoxanthin | -0.02152434138 | 0.0230126822 | 0.07510688278 | 1 |
| ergothioneine | -0.0130903173 | -0.004331751329 | 0.08447481424 | 1 |
| 4-acetamidophenol | 0.06218145268 | 0.02059019667 | 0.08447481424 | 1 |
| IF1AX_HUMAN_IF1AY_HUMAN | 0.1187914633 | 0.07511167789 | 0.1247717199 | 1 |
| dodecenedioate | -0.01671622674 | -0.003735426377 | 0.1422928156 | 1 |
| cysteine s-sulfate | 0.004629092434 | -0.005284178479 | 0.149800547 | 1 |
| MYOC_HUMAN | -0.02538097517 | -0.003411835789 | 0.1741093598 | 1 |
| IBP5_HUMAN | -0.01111721085 | 0.003172535683 | 0.1918552587 | 1 |

| | | | | |
|---|---|---|---|---|
| **cis-3,4-methyleneheptanoylcarnitine** | -0.03117731868 | 0.002802896031 | 0.2059999892 | 1 |
| **PLMN_HUMAN** | -0.01285262518 | -0.00272200703 | 0.2108768051 | 1 |
| **trans-3,4-methyleneheptanoate** | -0.01572439032 | -0.001259150751 | 0.2208761433 | 1 |
| **LYSC_HUMAN** | -0.01071029253 | -0.005924461983 | 0.2259995395 | 1 |
| **PON1_HUMAN** | -0.02415664192 | 0.006106702842 | 0.2259995395 | 1 |
| **arachidoylcarnitine** | 0.03208989317 | 0.008411536511 | 0.231205966 | 1 |
| **GRN_HUMAN** | -0.007761660016 | -0.001692921048 | 0.2364958031 | 1 |
| **salicylate** | -0.0318091145 | -0.01288575272 | 0.2418694073 | 1 |
| **cystine** | 0.01731686939 | -0.0000964707368 | 0.2528692218 | 1 |
| **10-undecenoate** | -0.01387633951 | -0.004503585043 | 0.264207768 | 1 |
| **3-hydroxydodecanedioate*** | -0.007490774196 | -0.0002052763238 | 0.264207768 | 1 |
| **H4_HUMAN** | -0.01529916186 | -0.001277547441 | 0.2758869914 | 1 |
| **ribitol** | 0.003925455011 | 0.001988467737 | 0.2818548481 | 1 |
| **THRB_HUMAN** | -0.004293647915 | 0.01420904353 | 0.2818548481 | 1 |
| **glycosyl-N-behenoyl-sphingadienine** | -0.007524278202 | 0.003520880956 | 0.2940477923 | 1 |
| **eicosenedioate** | -0.01847332257 | -0.005341254794 | 0.2940477923 | 1 |
| **F10A1_HUMAN** | 0.03744465736 | 0.01937536847 | 0.2940477923 | 1 |
| **ITIH3_HUMAN** | -0.02623582386 | -0.01072724112 | 0.2940477923 | 1 |
| **2-methylcitrate/homocitrate** | -0.02813440407 | -0.0115981768 | 0.3002730913 | 1 |
| **indolelactate** | 0.05467739248 | 0.03169813047 | 0.3065843675 | 1 |
| **gamma-glutamyltyrosine** | -0.007694903346 | -0.003614215427 | 0.3065843675 | 1 |
| **BMP1_HUMAN** | -0.006411807805 | 0.002223433066 | 0.3065843675 | 1 |
| **2-oxindole-3-acetate** | -0.01721888668 | -0.01043444506 | 0.3129816539 | 1 |
| **PON1_HUMAN** | -0.01280620599 | -0.004620669967 | 0.3194649531 | 1 |
| **1-palmitoyl-2-docosahexaenoyl-GPE** | -0.01992933466 | -0.007940208012 | 0.3260342379 | 1 |
| **1-margaroyl-GPE** | -0.01835847447 | -0.0042041271 | 0.33268945 | 1 |
| **GPX3_HUMAN** | -0.02186469303 | 0.007206006385 | 0.33268945 | 1 |
| **taurocholenate sulfate*** | -0.02777293117 | -0.006527503121 | 0.3394305005 | 1 |
| **sphingomyelin** | -0.03574589937 | -0.01884072551 | 0.3394305005 | 1 |
| **IBP2_HUMAN** | -0.01913602535 | -0.01118384801 | 0.3394305005 | 1 |
| **LYSC_HUMAN** | -0.01128067127 | -0.002719429248 | 0.3394305005 | 1 |
| **PGK1_HUMAN** | 0.02405111675 | 0.002856418465 | 0.3394305005 | 1 |
| **1-palmitoyl-2-eicosapentaenoyl-GPE** | -0.01626459169 | 0.00573745307 | 0.3462572689 | 1 |
| **APOF_HUMAN** | -0.02655432662 | -0.00004982877665 | 0.353169603 | 1 |
| **sphingomyelin** | -0.03815752943 | -0.02805092348 | 0.3601673188 | 1 |
| **PLMN_HUMAN** | -0.003705730379 | 0.0009144782042 | 0.3601673188 | 1 |
| **sebacate** | -0.0229189925 | -0.001349728651 | 0.3672502 | 1 |

| | | | | |
|---|---|---|---|---|
| **N-delta-acetylornithine** | -0.02014657162 | -0.0224597498 | 0.3672502 | 1 |
| **phenylalanine** | -0.002559750217 | 0.0004029856423 | 0.3672502 | 1 |
| **CSPG2_HUMAN** | -0.03220528088 | 0.0009592195763 | 0.3744179979 | 1 |
| **3beta-hydroxy-5-cholenoate** | 0.01021087053 | 0.008582375557 | 0.3816704308 | 1 |
| **PA2GA_HUMAN** | -0.0460750291 | -0.002356349687 | 0.3890071844 | 1 |
| **ALBU_HUMAN** | 0.01245659717 | 0.003448553838 | 0.3890071844 | 1 |
| **FETUA_HUMAN** | -0.002544661916 | 0.0006495502442 | 0.3890071844 | 1 |
| **ERF1_HUMAN** | 0.02144751519 | 0.02131754026 | 0.3964279109 | 1 |
| **CBPB2_HUMAN** | -0.01936306127 | -0.008518487733 | 0.3964279109 | 1 |
| **N-acetylglucosamine/N-acetylgalactosamine** | -0.01706500515 | -0.01049056794 | 0.4039322295 | 1 |
| **carnitine** | -0.01122231587 | -0.002699078344 | 0.4115197256 | 1 |
| **2-aminoheptanoate** | -0.01217482591 | 0.004210227323 | 0.4115197256 | 1 |
| **carotene diol** | -0.0005344990404 | 0.01392051032 | 0.4115197256 | 1 |
| **BMP1_HUMAN** | 0.0084700422 | -0.001926858747 | 0.4115197256 | 1 |
| **phenylacetylglycine** | 0.009827523587 | 0.00894449021 | 0.4191899511 | 1 |
| **5alpha-pregnan-3beta,20beta-diol monosulfate** | -0.03808263306 | -0.0005983687951 | 0.4191899511 | 1 |
| **FETUA_HUMAN** | -0.00114524397 | 0.0002363069786 | 0.4191899511 | 1 |
| **HGFA_HUMAN** | -0.009020385942 | -0.009849948578 | 0.4191899511 | 1 |
| **ITIH3_HUMAN** | -0.001335996789 | 0.001722852853 | 0.4191899511 | 1 |
| **APOF_HUMAN** | -0.02921141817 | -0.01614905947 | 0.4191899511 | 1 |
| **CSPG2_HUMAN** | -0.02518437442 | -0.004033361175 | 0.4269424242 | 1 |
| **PON1_HUMAN** | -0.001240246558 | 0.01396261538 | 0.4269424242 | 1 |
| **LYSC_HUMAN** | -0.01248833925 | -0.01008061005 | 0.4347766292 | 1 |
| **beta-citrylglutamate** | -0.001105186113 | -0.003364409352 | 0.4426920165 | 1 |
| **p-cresol glucuronide*** | 0.02011590208 | -0.01430107298 | 0.4426920165 | 1 |
| **IBP2_HUMAN** | -0.02289854482 | -0.007659849809 | 0.4426920165 | 1 |
| **RNAS4_HUMAN** | -0.003559472038 | -0.002767522607 | 0.4426920165 | 1 |
| **4-guanidinobutanoate** | 0.04703870818 | 0.04203380159 | 0.4506880026 | 1 |
| **uridine** | -0.0211908296 | 0.005462500276 | 0.4587639698 | 1 |
| **dihydroorotate** | 0.007926341108 | -0.0008497823571 | 0.4587639698 | 1 |
| **Total_cholesterol** | -4.624280001 | -3.83459081 | 0.4587639698 | 1 |
| **PGK1_HUMAN** | 0.006871433023 | -0.00120142666 | 0.4587639698 | 1 |
| **CFAD_HUMAN** | -0.005989975873 | 0.009261831939 | 0.4587639698 | 1 |
| **PON1_HUMAN** | -0.01883224515 | -0.008347055662 | 0.4669192668 | 1 |
| **glycochenodeoxycholate 3-sulfate** | 0.006168702684 | 0.003979502599 | 0.4669192668 | 1 |
| **1-palmitoyl-2-arachidonoyl-GPE** | -0.01896009868 | 0.001702559206 | 0.4751532079 | 1 |

| | | | | |
|---|---|---|---|---|
| **docosahexaenoylcarnitine** | -0.006825679693 | -0.001072570645 | 0.4751532079 | 1 |
| **PON1_HUMAN** | -0.02925679353 | -0.02201158276 | 0.4751532079 | 1 |
| **H2B1_HUMAN** | -0.01374356896 | -0.00489090163 | 0.4751532079 | 1 |
| **ITIH3_HUMAN** | -0.01703477983 | -0.007921790686 | 0.4751532079 | 1 |
| **FBLN3_HUMAN** | -0.009602824991 | 0.006279682175 | 0.4751532079 | 1 |
| **MXRA5_HUMAN** | 0.003772095216 | 0.01861752144 | 0.4834650739 | 1 |
| **RNAS1_HUMAN** | -0.0120334541 | -0.002327662933 | 0.4834650739 | 1 |
| **CBPB2_HUMAN** | -0.003975206171 | -0.01876349819 | 0.4918541113 | 1 |
| **CMPF** | -0.0459991495 | 0.0003921826782 | 0.4918541113 | 1 |
| **N-acetylmethionine** | -0.007564390664 | 0.001117553943 | 0.4918541113 | 1 |
| **CCL14_HUMAN** | -0.01614636665 | -0.004417319577 | 0.4918541113 | 1 |
| **PON1_HUMAN** | -0.008115168108 | -0.006001703325 | 0.4918541113 | 1 |
| **vanillylmandelate** | -0.02712158899 | -0.008818375362 | 0.5003195333 | 1 |
| **F10A1_HUMAN** | 0.05436068896 | 0.03974745911 | 0.5088605191 | 1 |
| **myristoylcarnitine** | -0.01267422048 | -0.01062516238 | 0.5088605191 | 1 |
| **1-linoleoyl-2-arachidonoyl-GPC** | 0.01325855824 | 0.006246492012 | 0.5088605191 | 1 |
| **3-4-hydroxyphenyl-lactate** | 0.01799806996 | 0.007420691025 | 0.5088605191 | 1 |
| **CRDL1-4_HUMAN** | -0.01187393316 | -0.003257398816 | 0.5088605191 | 1 |
| **SLPI_HUMAN** | 0.01049731583 | 0.003413850279 | 0.5088605191 | 1 |
| **HAP28_HUMAN** | 0.004741626397 | -0.001301230226 | 0.5088605191 | 1 |
| **N-acetylalliin** | -0.01239847646 | 0.001629284894 | 0.5174762146 | 1 |
| **N-carbamoylalanine** | 0.03998421759 | 0.03674482344 | 0.5174762146 | 1 |
| **carnitine of C10H14O2** | -0.003694305313 | 0.001478593931 | 0.5174762146 | 1 |
| **sphingomyelin** | -0.09544509045 | -0.1153190864 | 0.5261657322 | 1 |
| **1-1-enyl-palmitoyl-2-linoleoyl-GPC** | -0.00136050132 | -0.0002234433661 | 0.5261657322 | 1 |
| **NOTUM_HUMAN** | -0.02566314436 | 0.001418268075 | 0.5349281514 | 1 |
| **iminodiacetate** | 0.002740393609 | 0.00488869225 | 0.5349281514 | 1 |
| **tyramine O-sulfate** | 0.005849932331 | -0.003944932793 | 0.5349281514 | 1 |
| **4-hydroxyphenylacetylglutamine** | -0.0372157041 | -0.004803592133 | 0.5349281514 | 1 |
| **3-hydroxyoleoylcarnitine** | -0.01313964753 | -0.002863556827 | 0.5349281514 | 1 |
| **methionine sulfone** | 0.01217258693 | 0.01350053248 | 0.5437625185 | 1 |
| **PCOC1_HUMAN** | -0.001422925623 | 0.001689702161 | 0.5437625185 | 1 |
| **CO6A3_HUMAN** | 0.02037644764 | 0.0001615236839 | 0.5437625185 | 1 |
| **3-methylcytidine** | -0.00172083939 | 0.005256916525 | 0.5526678473 | 1 |
| **tryptophan betaine** | -0.01798027853 | -0.006821874704 | 0.5526678473 | 1 |
| **behenoyl dihydrosphingomyelin** | -0.01179689509 | -0.002219379243 | 0.5526678473 | 1 |
| **citrulline** | 0.0105840359 | -0.0003094816966 | 0.5526678473 | 1 |

| | | | | |
|---|---|---|---|---|
| **gentisate** | 0.003684641297 | 0.002568870161 | 0.5616431191 | 1 |
| **21-hydroxypregnenolone disulfate** | -0.003942363423 | 0.02211962529 | 0.5616431191 | 1 |
| **PON1_HUMAN** | 0.003102532734 | 0.007510234529 | 0.5616431191 | 1 |
| **COIA1_HUMAN** | -0.001246887732 | 0.001142410826 | 0.5616431191 | 1 |
| **quinate** | -0.02108596122 | 0.01053968786 | 0.5706872829 | 1 |
| **5-hydroxyhexanoate** | -0.01307891992 | -0.003204525244 | 0.5706872829 | 1 |
| **CCL16_HUMAN** | -0.009484502955 | -0.00556382667 | 0.5706872829 | 1 |
| **2-hydroxy-4-methylthio-butanoic acid** | 0.01285385702 | 0.004055667327 | 0.5797992561 | 1 |
| **branched-chain** | -0.001676735102 | -0.0000485145252<br>4 | 0.5797992561 | 1 |
| **PCOC1_HUMAN** | -0.002400245553 | -0.005958733023 | 0.5797992561 | 1 |
| **PON1_HUMAN** | -0.00115005082 | -0.01126286857 | 0.5889779245 | 1 |
| **Smoking_PackYears** | -0.3773867767 | -0.5421617508 | 0.5889779245 | 1 |
| **PCOC1_HUMAN** | -0.001512936817 | -0.003801401276 | 0.5889779245 | 1 |
| **BUN** | -0.3489588783 | 0.4993756357 | 0.5982221427 | 1 |
| **phenylacetylcarnitine** | -0.02282197684 | -0.005116232616 | 0.5982221427 | 1 |
| **glucuronide of piperine metabolite** | 0.02558827999 | 0.03271228408 | 0.5982221427 | 1 |
| **serotonin** | -0.004393564183 | -0.0107027819 | 0.5982221427 | 1 |
| **IBP6_HUMAN** | -0.01292064006 | -0.005003766601 | 0.5982221427 | 1 |
| **N-methylpipecolate** | 0.01213120686 | -0.01095523588 | 0.6075307345 | 1 |
| **malonate** | 0.004218631529 | -0.001123583677 | 0.6075307345 | 1 |
| **sucrose** | -0.01088769964 | -0.01954424283 | 0.6075307345 | 1 |
| **Lipid_HDL** | -0.2113849259 | -0.8346921048 | 0.6075307345 | 1 |
| **RNAS1_HUMAN** | -0.02128459742 | -0.003250287967 | 0.6075307345 | 1 |
| **1-pentadecanoyl-GPC** | 0.01123602682 | 0.005298430012 | 0.6169024936 | 1 |
| **hypoxanthine** | -0.00397493862 | 0.003530041456 | 0.6169024936 | 1 |
| **HbA1c** | -0.001094187447 | -0.05222220277 | 0.6169024936 | 1 |
| **PCOC1_HUMAN** | -0.009117782514 | -0.009536472138 | 0.6169024936 | 1 |
| **7-hydroxyindole sulfate** | -0.006989631676 | 0.001260323411 | 0.6263361834 | 1 |
| **carotene diol** | -0.00248095836 | 0.01327366633 | 0.6263361834 | 1 |
| **glycine conjugate of C10H14O2** | -0.007344401229 | 0.006480241886 | 0.6263361834 | 1 |
| **androsterone glucuronide** | 0.01254415169 | 0.007872787978 | 0.6358305382 | 1 |
| **lyxonate** | -0.02417287576 | -0.001971750824 | 0.6358305382 | 1 |
| **glutamine conjugate of C7H12O2*** | -0.02100671585 | 0.003154501771 | 0.6358305382 | 1 |
| **retinol_vitamin A** | 0.01235473448 | -0.008253217224 | 0.6358305382 | 1 |
| **glutamate** | 0.03898497633 | 0.0297465838 | 0.6358305382 | 1 |
| **SLPI_HUMAN** | 0.001805903641 | 0.003051814161 | 0.6358305382 | 1 |
| **1-palmitoyl-GPC** | -0.004301871921 | 0.004407061717 | 0.6453842629 | 1 |

| | | | | |
|---|---|---|---|---|
| **N-oleoyltaurine** | 0.009295520558 | -0.005011623514 | 0.6453842629 | 1 |
| **carotene diol** | -0.01064540802 | -0.003231234442 | 0.6453842629 | 1 |
| **hydroxyasparagine**** | -0.04546731167 | -0.01681801525 | 0.6453842629 | 1 |
| **BMP1_HUMAN** | -0.0009616211434 | 0.001101835292 | 0.6453842629 | 1 |
| **2-hydroxyglutarate** | 0.003779134298 | -0.0002734598228 | 0.6549960342 | 1 |
| **4-hydroxyglutamate** | -0.004235470846 | 0.009525204186 | 0.6646645006 | 1 |
| **behenoyl sphingomyelin** | -0.007128648829 | -0.0118510328 | 0.6646645006 | 1 |
| **pantothenate** | 0.009301085151 | 0.003499720888 | 0.6646645006 | 1 |
| **BMP1_HUMAN** | 0.00527825787 | -0.001151059982 | 0.6646645006 | 1 |
| **PCOC1_HUMAN** | -0.001824963364 | -0.01341448223 | 0.6646645006 | 1 |
| **BMP1_HUMAN_TLL1_HUMAN** | 0.008279509358 | 0.009116943316 | 0.6646645006 | 1 |
| **21-hydroxypregnenolone monosulfate** | 0.02047844821 | 0.02505541364 | 0.6743882834 | 1 |
| **3-methoxytyramine sulfate** | -0.0004267236607 | 0.001683939357 | 0.6743882834 | 1 |
| **N,N,N-trimethyl-5-aminovalerate** | 0.001942971889 | 0.0003579691883 | 0.6743882834 | 1 |
| **3,5-dichloro-2,6-dihydroxybenzoic acid** | 0.00514940974 | -0.0006568930813 | 0.6743882834 | 1 |
| **spermidine** | -0.001055948049 | 0.0004256326116 | 0.6743882834 | 1 |
| **adenosine** | 0.005796652126 | -0.000718190967 | 0.6743882834 | 1 |
| **PON1_HUMAN** | 0.001434378314 | 0.002342310376 | 0.6743882834 | 1 |
| **salicyluric glucuronide*** | -0.02802135372 | -0.04918217544 | 0.6841659768 | 1 |
| **5-methyluridine_ribothymidine** | -0.004379219515 | -0.005729769515 | 0.6841659768 | 1 |
| **N,N-dimethyl-5-aminovalerate** | -0.01225383669 | -0.007222532822 | 0.6841659768 | 1 |
| **indole-3-carboxylate** | -0.003843265705 | 0.004234424085 | 0.6939961488 | 1 |
| **1,2-dilinoleoyl-GPC** | 0.004504610847 | 0.006583628895 | 0.6939961488 | 1 |
| **hexanoylcarnitine** | -0.02796705242 | -0.005658845869 | 0.7038773417 | 1 |
| **1-stearoyl-2-adrenoyl-GPC** | 0.01036727773 | 0.004905264399 | 0.7038773417 | 1 |
| **1,2-dipalmitoyl-GPC** | -0.000297902569 | -0.002856682566 | 0.7138080727 | 1 |
| **N2-acetyllysine** | 0.004992108324 | 0.0210842406 | 0.7138080727 | 1 |
| **N-linoleoyltaurine*** | 0.003934595201 | 0.009501637368 | 0.7138080727 | 1 |
| **2,4-di-tert-butylphenol** | -0.01311637984 | -0.00690386642 | 0.7138080727 | 1 |
| **urate** | 0.04598383487 | 0.03331258591 | 0.7138080727 | 1 |
| **Lipid_LDL** | -1.651187072 | -1.146644672 | 0.7138080727 | 1 |
| **androsterone sulfate** | 0.02676018456 | -0.006322034763 | 0.7237868346 | 1 |
| **N-acetylglutamine** | -0.005589500902 | -0.001707110232 | 0.7237868346 | 1 |
| **3-ureidopropionate** | 0.01204562577 | -0.0007545460689 | 0.7237868346 | 1 |
| **beta-hydroxyisovalerate** | 0.02787143188 | 0.01442080084 | 0.7237868346 | 1 |
| **leucine** | 0.008253622193 | 0.01104221869 | 0.7237868346 | 1 |

| | | | | |
|---|---|---|---|---|
| **H4_HUMAN** | -0.0113587229 | 0.009100303614 | 0.7237868346 | 1 |
| **4-methoxyphenol sulfate** | -0.02478983565 | -0.01084022182 | 0.7338120965 | 1 |
| **guanidinosuccinate** | 0.01181040988 | -0.0008179752311 | 0.7338120965 | 1 |
| **N-acetyl-3-methylhistidine*** | 0.03452016005 | 0.004126826187 | 0.7338120965 | 1 |
| **hydroxybutyrylcarnitine** | -0.009805178995 | -0.001158050556 | 0.7338120965 | 1 |
| **malate** | 0.0002244056781 | -0.002880515035 | 0.7338120965 | 1 |
| **urea** | 0.003363306669 | -0.0002111459859 | 0.7338120965 | 1 |
| **CIRBP_HUMAN** | 0.005170159797 | -0.01428493104 | 0.7338120965 | 1 |
| **1,5-anhydroglucitol** | -0.002411641815 | -0.001954732556 | 0.7438823044 | 1 |
| **phenylacetylglutamine** | -0.000811366007 | 0.002898187241 | 0.7438823044 | 1 |
| **androsterone glucuronide** | 0.03994957598 | -0.005667850182 | 0.7438823044 | 1 |
| **argininate*** | 0.01454677278 | 0.01095808085 | 0.7438823044 | 1 |
| **succinimide** | 0.009684818479 | 0.02629895971 | 0.7438823044 | 1 |
| **2-hydroxyphytanate*** | -0.001163002157 | -0.0003021092455 | 0.7438823044 | 1 |
| **trans-2-hexenoylglycine** | -0.002921676589 | -0.003093324553 | 0.7438823044 | 1 |
| **N-acetyl-2-aminooctanoate*** | 0.01768782038 | 0.01202556645 | 0.7438823044 | 1 |
| **creatinine** | 0.04581507263 | 0.03460854335 | 0.7438823044 | 1 |
| **ornithine** | -0.002709040851 | 0.0003870842184 | 0.7438823044 | 1 |
| **IBP6_HUMAN** | -0.001736859328 | 0.00000340195048 | 0.7438823044 | 1 |
| **11-ketoetiocholanolone glucuronide** | -0.003488207661 | -0.002263474378 | 0.7539958819 | 1 |
| **1-stearoyl-2-dihomo-linolenoyl-GPC** | -0.01080973943 | 0.0004904677984 | 0.7539958819 | 1 |
| **catechol glucuronide** | -0.006638382658 | -0.001979816122 | 0.7539958819 | 1 |
| **levulinoylcarnitine** | -0.03232038881 | -0.02298504915 | 0.7539958819 | 1 |
| **Red_Cell_Dist_Width** | -0.02768352386 | 0.04248710144 | 0.764151231 | 1 |
| **N-acetyl-cadaverine** | 0.01112135523 | -0.0004995026283 | 0.764151231 | 1 |
| **N-stearoyl-sphingosine** | 0.008081239299 | -0.005976290188 | 0.764151231 | 1 |
| **FVC** | 0.03152089063 | 0.1199313496 | 0.764151231 | 1 |
| **Total_Bilirubin** | 0.01127231823 | 0.02543383211 | 0.764151231 | 1 |
| **H4_HUMAN** | -0.001871166025 | 0.007065705233 | 0.764151231 | 1 |
| **IBP6_HUMAN** | 0.00567077806 | -0.004434789513 | 0.764151231 | 1 |
| **CBPB2_HUMAN** | -0.00522181652 | -0.0034306589 | 0.764151231 | 1 |
| **gamma-glutamylglycine** | 0.005280937564 | -0.005127218066 | 0.7743467329 | 1 |
| **glutamine_degradant*** | 0.03157239674 | 0.04505798109 | 0.7743467329 | 1 |
| **chiro-inositol** | 0.002812665617 | -0.001241232021 | 0.7743467329 | 1 |
| **cinnamoylglycine** | -0.01842136169 | 0.006415178086 | 0.7743467329 | 1 |
| **1-methyl-5-imidazoleacetate** | -0.002779697985 | -0.0007840293948 | 0.7743467329 | 1 |
| **1-margaroyl-2-arachidonoyl-GPC** | 0.006375764623 | -0.00434094222 | 0.7743467329 | 1 |

| | | | | |
|---|---|---|---|---|
| **S-carboxyethylcysteine** | -0.004932428975 | -0.003458015037 | 0.7743467329 | 1 |
| **N1-methyladenosine** | -0.006147127738 | 0.003777355238 | 0.7743467329 | 1 |
| **alpha-ketoglutarate** | -0.004347028003 | 0.001011221059 | 0.7743467329 | 1 |
| **Creatinine** | 0.01790761186 | 0.05981818206 | 0.7845807485 | 1 |
| **N-acetylglutamate** | 0.002605341386 | 0.003398197013 | 0.7845807485 | 1 |
| **dimethyl sulfone** | 0.007322108301 | -0.008238885098 | 0.7845807485 | 1 |
| **thyroxine** | -0.01092529013 | -0.0030217041 | 0.7845807485 | 1 |
| **riboflavin_vitamin B2** | 0.002714678224 | 0.0003114881634 | 0.7845807485 | 1 |
| **mannose** | 0.0000452245773 4 | -0.008788299515 | 0.7845807485 | 1 |
| **IBP2_HUMAN** | -0.000774833693 | -0.01731543688 | 0.7845807485 | 1 |
| **dehydroepiandrosterone sulfate** | 0.007183633549 | -0.01714418736 | 0.7948516193 | 1 |
| **galactonate** | 0.008612631455 | -0.003267978425 | 0.7948516193 | 1 |
| **3-methyladipate** | -0.02447109347 | -0.02068788117 | 0.7948516193 | 1 |
| **6-oxopiperidine-2-carboxylate** | 0.0005515196986 | -0.000091081602 | 0.7948516193 | 1 |
| **1-pentadecanoyl-2-linoleoyl-GPC** | 0.001947939006 | 0.0008825952121 | 0.7948516193 | 1 |
| **undecenoylcarnitine** | 0.000201498693 | 0.003840921933 | 0.7948516193 | 1 |
| **acetoacetate** | -0.01205742786 | -0.0004589688844 | 0.7948516193 | 1 |
| **PCOC1_HUMAN** | -0.01174933553 | -0.001244751101 | 0.7948516193 | 1 |
| **CBPB2_HUMAN** | -0.004494451143 | -0.01641263144 | 0.8051576685 | 1 |
| **2-hydroxysebacate** | -0.009769557297 | -0.005490642275 | 0.8051576685 | 1 |
| **3-hydroxyoctanoylcarnitine** | -0.005041560113 | 0.006622790813 | 0.8051576685 | 1 |
| **alpha-tocopherol** | -0.004557641039 | 0.002599479501 | 0.8051576685 | 1 |
| **CCL18_HUMAN** | -0.003001495916 | 0.0001111969152 | 0.8051576685 | 1 |
| **sphinganine-1-phosphate** | -0.009261750683 | -0.01428088572 | 0.8154972013 | 1 |
| **2-hydroxybutyrate/2-hydroxyisobutyr ate** | 0.0009706868634 | 0.004240879096 | 0.8154972013 | 1 |
| **1-myristoyl-2-arachidonoyl-GPC** | -0.006195463345 | -0.01043604299 | 0.8154972013 | 1 |
| **guanidinoacetate** | 0.01012044055 | 0.01336450462 | 0.8154972013 | 1 |
| **MMP19_HUMAN** | -0.02653603468 | 0.001681108828 | 0.8154972013 | 1 |
| **MGP_HUMAN** | -0.001503776746 | -0.005054307908 | 0.8154972013 | 1 |
| **homoarginine** | 0.006237072837 | 0.003347174475 | 0.8258685061 | 1 |
| **1-docosahexaenoylglycerol** | -0.007467364725 | 0.004369252343 | 0.8258685061 | 1 |
| **omeprazole** | 0.008992665199 | 0.001327141259 | 0.8258685061 | 1 |
| **ferulic acid 4-sulfate** | -0.003579196029 | -0.004116613562 | 0.8258685061 | 1 |
| **lithocholic acid sulfate** | -0.001507843427 | -0.02491803623 | 0.8258685061 | 1 |
| **menthol glucuronide** | 0.002054607562 | 0.00626310139 | 0.8258685061 | 1 |
| **FEV1** | 0.01263564479 | -0.0003738401406 | 0.8258685061 | 1 |

| | | | | |
|---|---|---|---|---|
| **androstenediol disulfate** | 0.0254844542 | 0.02143930472 | 0.8362698549 | 1 |
| **ximenoylcarnitine** | 0.0104466661 | 0.001750193861 | 0.8362698549 | 1 |
| **glucuronide of C12H22O4** | 0.004362244735 | -0.003100376177 | 0.8362698549 | 1 |
| **glucuronide of C10H18O2** | 0.01690139319 | -0.01185645254 | 0.8362698549 | 1 |
| **dihydroferulic acid sulfate** | 0.00546823586 | -0.007871151349 | 0.8362698549 | 1 |
| **cystathionine** | -0.001756235618 | -0.0119903382 | 0.8362698549 | 1 |
| **gluconate** | 0.01308601975 | 0.004552692306 | 0.8362698549 | 1 |
| **IBP2_HUMAN** | -0.01788424862 | 0.0006890374773 | 0.8362698549 | 1 |
| **IBP2_HUMAN** | -0.01219100786 | -0.01232208865 | 0.8362698549 | 1 |
| **AMBP_HUMAN** | -0.000221366484 | -0.002371933696 | 0.8362698549 | 1 |
| **3-methylxanthine** | 0.01780127325 | 0.01388676794 | 0.8466995048 | 1 |
| **lactosyl-N-palmitoyl-sphingosine** | 0.0009454570552 | 0.01005823526 | 0.8466995048 | 1 |
| **isocitric lactone** | -0.0006431301146 | -0.01121163584 | 0.8466995048 | 1 |
| **picolinate** | 0.008181046817 | -0.005672628929 | 0.8466995048 | 1 |
| **estrone 3-sulfate** | 0.01789460219 | -0.004930396075 | 0.8466995048 | 1 |
| **methylsuccinate** | 0.009148350372 | 0.008826990704 | 0.8466995048 | 1 |
| **lactose** | -0.0180915841 | -0.01717273864 | 0.8466995048 | 1 |
| **Direct_Bilirubin** | 0.00346949145 | -0.002952313422 | 0.8466995048 | 1 |
| **CYTC_HUMAN** | 0.004358411739 | 0.006052196772 | 0.8466995048 | 1 |
| **PCOC1_HUMAN** | -0.001657621409 | -0.003863431324 | 0.8466995048 | 1 |
| **FETUA_HUMAN** | -0.00166548027 | -0.002959093398 | 0.8466995048 | 1 |
| **indoleacetylglutamine** | -0.01210635073 | -0.01258944035 | 0.8571556983 | 1 |
| **suberoylcarnitine** | -0.006500858173 | -0.003736511793 | 0.8571556983 | 1 |
| **3-hydroxyphenylacetoylglutamine** | 0.004760548423 | -0.0007677129168 | 0.8571556983 | 1 |
| **3-hydroxyhippurate sulfate** | -0.008923533731 | -0.004445531048 | 0.8571556983 | 1 |
| **picolinoylglycine** | -0.01636589977 | -0.008805926599 | 0.8571556983 | 1 |
| **3-methoxytyrosine** | 0.001078768735 | -0.00101036535 | 0.8571556983 | 1 |
| **uracil** | -0.0006571386716 | 0.008711979628 | 0.8571556983 | 1 |
| **caffeine** | -0.004290710557 | 0.005980318959 | 0.8571556983 | 1 |
| **BMP1_HUMAN** | 0.001998726076 | 0.007976298157 | 0.8571556983 | 1 |
| **CRP** | -0.5716600154 | -0.8795429204 | 0.8676366644 | 1 |
| **Triglyceride** | 6.929036211 | 8.419515568 | 0.8676366644 | 1 |
| **1-methylguanidine** | -0.001874769613 | 0.009839243521 | 0.8676366644 | 1 |
| **2-methoxyhydroquinone sulfate** | 0.002404068687 | -0.005701643001 | 0.8676366644 | 1 |
| **xanthine** | -0.0007086834362 | 0.004437497101 | 0.8676366644 | 1 |
| **BMI** | -0.398626512 | -0.4265069556 | 0.8781406193 | 1 |
| **Mean_Corpus_Vol** | -0.005140046971 | -0.07462663989 | 0.8781406193 | 1 |

| | | | | |
|---|---|---|---|---|
| **gamma-glutamylphenylalanine** | -0.006296186425 | -0.008285952429 | 0.8781406193 | 1 |
| **N-acetyl-1-methylhistidine*** | 0.01217369022 | 0.001264037307 | 0.8781406193 | 1 |
| **IBP2_HUMAN** | 0.008567214547 | -0.01127486066 | 0.8781406193 | 1 |
| **Hematocrit** | -0.02747061677 | 0.0117766837 | 0.8886657678 | 1 |
| **Hemoglobin** | 0.07655711043 | 0.02614875731 | 0.8886657678 | 1 |
| **lanthionine** | 0.0102448451 | -0.01082469032 | 0.8886657678 | 1 |
| **3-hydroxyadipate** | -0.002938183935 | -0.00271150523 | 0.8886657678 | 1 |
| **histidine** | -0.004385928722 | -0.005372601937 | 0.8886657678 | 1 |
| **PON1_HUMAN** | 0.002665147356 | -0.005966065352 | 0.8886657678 | 1 |
| **vanillactate** | -0.01035991041 | 0.01043643217 | 0.8992103035 | 1 |
| **citramalate** | -0.002211968243 | -0.002238572268 | 0.8992103035 | 1 |
| **1-palmitoyl-2-linoleoyl-GPC** | -0.0000724805510 6 | 0.0006533241959 | 0.8992103035 | 1 |
| **choline phosphate** | 0.005983431112 | 0.0006186251196 | 0.8992103035 | 1 |
| **Glucose** | 0.5460330662 | -0.3422180118 | 0.9097724102 | 1 |
| **threonate** | 0.00324543569 | 0.004827587657 | 0.9097724102 | 1 |
| **indolebutyrate** | -0.006053224109 | -0.008844893767 | 0.9097724102 | 1 |
| **mannonate*** | -0.01262400715 | -0.02350412979 | 0.9097724102 | 1 |
| **dopamine 4-sulfate** | 0.01279533244 | -0.009267382023 | 0.9097724102 | 1 |
| **methyl vanillate sulfate** | -0.0004848661104 | 0.0002661157931 | 0.9097724102 | 1 |
| **PCOC1_HUMAN** | -0.00341107377 | -0.000744108349 | 0.9097724102 | 1 |
| **IBP2_HUMAN** | 0.005669370068 | 0.001659763185 | 0.9097724102 | 1 |
| **IBP2_HUMAN** | 0.006907153976 | -0.008835295104 | 0.9203502627 | 1 |
| **acetylcarnitine** | 0.009398454927 | -0.01214770982 | 0.9203502627 | 1 |
| **furaneol sulfate** | 0.0000694288123 6 | 0.0007145409782 | 0.9203502627 | 1 |
| **decadienedioic acid** | -0.001261484486 | -0.00507642598 | 0.9203502627 | 1 |
| **ibuprofen** | -0.0001368705989 | -0.0006268682788 | 0.9203502627 | 1 |
| **Liver_ALB** | 0.002274890215 | 0.01445233155 | 0.9309420277 | 1 |
| **IGF1_HUMAN** | 0.005102323727 | -0.006883971718 | 0.9309420277 | 1 |
| **indoleacetate** | 0.006320481392 | 0.003035460761 | 0.9309420277 | 1 |
| **nicotinamide riboside** | -0.0001972355007 | -0.0006034880904 | 0.9309420277 | 1 |
| **deoxycarnitine** | 0.002142700224 | 0.01738635548 | 0.9309420277 | 1 |
| **dimethylarginine** | -0.005413716245 | -0.007170778027 | 0.9309420277 | 1 |
| **arabitol/xylitol** | -0.02949414595 | -0.01379063825 | 0.9309420277 | 1 |
| **6-bromotryptophan** | 0.01093570922 | 0.003640162619 | 0.9309420277 | 1 |
| **5-hydroxymethyl-2-furoylcarnitine*** | -0.00876184371 | -0.02709432722 | 0.9309420277 | 1 |
| **TLL1_HUMAN** | 0.009535319377 | 0.006381144214 | 0.9309420277 | 1 |

| | | | | |
|---|---|---|---|---|
| **hippurate** | -0.005484805353 | -0.01370285513 | 0.9415458647 | 1 |
| **octadecanedioylcarnitine** | -0.001920106126 | 0.01539395498 | 0.9415458647 | 1 |
| **3-hydroxyindolin-2-one sulfate** | -0.005583347423 | 0.002408696987 | 0.9415458647 | 1 |
| **cortisol** | -0.00276673417 | -0.008359807866 | 0.9415458647 | 1 |
| **tryptophan** | 0.0023194349 | 0.004385611456 | 0.9415458647 | 1 |
| **Liver_ALP** | -0.5202097593 | -0.3278945479 | 0.9415458647 | 1 |
| **PCOC1_HUMAN** | -0.005815337439 | -0.005311710092 | 0.9415458647 | 1 |
| **isobutyrylcarnitine** | 0.003634404402 | 0.007856259835 | 0.9521599269 | 1 |
| **N-acetyltyrosine** | 0.001499157086 | -0.001431785845 | 0.9521599269 | 1 |
| **vanillic acid glycine** | 0.003358128912 | 0.006815753867 | 0.9521599269 | 1 |
| **trans-4-hydroxyproline** | 0.001050541509 | 0.0002013719495 | 0.9521599269 | 1 |
| **cholesterol** | -0.005909076232 | -0.01143557609 | 0.9521599269 | 1 |
| **proline** | 0.002193867386 | 0.006171374381 | 0.9521599269 | 1 |
| **serine** | 0.007419858508 | 0.006975964788 | 0.9521599269 | 1 |
| **HGFA_HUMAN** | 0.0008048788769 | -0.0004928273272 | 0.9521599269 | 1 |
| **tiglylcarnitine** | 0.00890605837 | -0.005878639955 | 0.9627823626 | 1 |
| **5-galactosylhydroxy-lysine** | -0.003127925539 | -0.003650265848 | 0.9627823626 | 1 |
| **3-methoxycatechol sulfate** | 0.01158288223 | 0.004178811825 | 0.9627823626 | 1 |
| **N-acetyl-isoputreanine** | -0.00688828385 | -0.01325600412 | 0.9627823626 | 1 |
| **hypotaurine** | 0.007654375028 | -0.0009594021392 | 0.9627823626 | 1 |
| **MIME_HUMAN** | -0.009890481422 | 0.00003499864771 | 0.9627823626 | 1 |
| **2-acetamidophenol sulfate** | 0.02872467365 | 0.001920423765 | 0.9734113153 | 1 |
| **eicosenoylcarnitine** | 0.01246072377 | 0.004295425321 | 0.9734113153 | 1 |
| **nicotinamide** | 0.003200842443 | 0.006106192205 | 0.9734113153 | 1 |
| **BMP1_HUMAN** | 0.00141373686 | 0.0007753092263 | 0.9734113153 | 1 |
| **MXRA5_HUMAN** | 0.004216531297 | 0.00518780205 | 0.9734113153 | 1 |
| **IBP6_HUMAN** | -0.0119728877 | 0.002213881099 | 0.9734113153 | 1 |
| **xanthosine** | 0.01606745582 | 0.004446333901 | 0.9840449255 | 1 |
| **propionylcarnitine** | 0.01211956139 | 0.002399190106 | 0.9840449255 | 1 |
| **N-acetylcitrulline** | 0.01512303957 | 0.01039833419 | 0.9840449255 | 1 |
| **cyclo_gly-pro** | 0.001540823751 | -0.007246737006 | 0.9840449255 | 1 |
| **ethyl alpha-glucopyranoside** | -0.005334414086 | 0.01143897747 | 0.9840449255 | 1 |
| **3-amino-2-piperidone** | -0.01480953767 | -0.02453794322 | 0.9840449255 | 1 |
| **dimethylguanidino valeric acid** | 0.0030760457 | -0.003571011108 | 0.9840449255 | 1 |
| **arabinose** | 0.0009670106063 | 0.001658935387 | 0.9840449255 | 1 |
| **SLPI_HUMAN** | -0.001111681457 | -0.004872878765 | 0.9840449255 | 1 |
| **BMP1_HUMAN** | 0.00740404507 | 0.01128670942 | 0.9840449255 | 1 |

| | | | | |
|---|---|---|---|---|
| **PCOC1_HUMAN** | 0.003017403408 | 0.004182820073 | 0.9840449255 | 1 |
| **pregnanediol-3-glucuronide** | 0.009017756051 | 0.0004788910054 | 0.9946813312 | 1 |
| **alpha-CMBHC glucuronide** | -0.01787111475 | -0.008056942645 | 0.9946813312 | 1 |
| **stearoyl-arachidonoyl-glycerol** | 0.0007911031877 | 0.001003212236 | 0.9946813312 | 1 |
| **cytosine** | 0.006018701057 | 0.004673232414 | 0.9946813312 | 1 |
| **IBP2_HUMAN** | -0.0001295536412 | -0.00359491755 | 0.9946813312 | 1 |
| **CRDL1-4_HUMAN** | 0.005410291199 | 0.008543464954 | 0.9946813312 | 1 |
| **succinylcarnitine** | -0.003704064056 | 0.001034480478 | 0.9946813312 | 1 |
| **linoleoylcarnitine** | 0.01657949941 | -0.001438207362 | 0.9946813312 | 1 |
| **N-acetyl-S-allyl-cysteine** | -0.005829258957 | -0.0007050492053 | 0.9946813312 | 1 |
| **phenylacetylglutamate** | -0.001955216825 | -0.004953423538 | 0.9946813312 | 1 |
| **2,3-dihydroxy-2-methylbutyrate** | -0.01204520374 | -0.01996415325 | 0.9946813312 | 1 |
| **homovanillate** | -0.003007654632 | -0.004217732784 | 0.9946813312 | 1 |
| **CCL14_HUMAN** | -0.006253210793 | -0.006710378206 | 0.9946813312 | 1 |
| **PCOC1_HUMAN** | 0.00571273161 | 0.001057456291 | 0.9946813312 | 1 |
| **3-hydroxy-2-ethylpropionate** | -0.006379518365 | -0.01231421488 | 1 | 1 |
| **N1-methyl-2-pyridone-5-carboxamide** | -0.01157705713 | -0.01546103826 | 1 | 1 |
| **acetylspermidine** | 0.01174885454 | -0.0004996917207 | 1 | 1 |
| **3-hydroxybutyroylglycine**** | -0.01666987788 | -0.002321167331 | 1 | 1 |
| **glycine conjugate of C10H12O2*** | 0.001100153416 | 0.01073627757 | 1 | 1 |

**Supplementary Table 2. Marioni protein biomarker analysis between baseline and 12 months.** The first column details all Marioni proteins assessed between baseline and 12 months. The following columns (columns 2 and 3) report the predicted marginal mean values at each of the time points. These values were adjusted for confounding factors technical variability such as age, and array type.The p-value column depicts the p-value of the Wilcoxon-rank sum test performed between baseline and 12 months. The final column depicts the adjusted p-value by using the false discovery rate (FDR) adjustment.

| | **Mean** | | **0 vs 12 months** | |
|---|---|---|---|---|
| | **0 months** | **12 months** | **p-value** | **FDR** |
| **HGFA** | -0.004203092086 | -0.00002423180198 | 0.006633638129 | 0.776135661 |
| **CCL25.C.C** | 0.002364087425 | -0.0002437389103 | 0.04517312702 | 0.9989683987 |
| **Smoking** | 0.05098585389 | -0.05589807382 | 0.08928440953 | 0.9989683987 |
| **Complement.C5a** | -0.002406946082 | -0.0005270524153 | 0.09389576275 | 0.9989683987 |
| **CCL11** | 0.001569555276 | 0.0000639118954 | 0.09707473754 | 0.9989683987 |
| **CCL17** | 0.002145944585 | -0.002608575736 | 0.127691418 | 0.9989683987 |
| **LFT** | -0.001969481142 | -0.0005748644109 | 0.127691418 | 0.9989683987 |

| | | | | |
|---|---|---|---|---|
| **G.CSF** | 0.00002570528647 | -0.00196911118 | 0.1778664331 | 0.9989683987 |
| **SLITRK5** | -0.002253284984 | 0.002162560076 | 0.1910861944 | 0.9989683987 |
| **Complement.c9** | -0.003249777039 | 0.002421814867 | 0.1993575334 | 0.9989683987 |
| **CD6** | 0.000539745659 | -0.002415556568 | 0.2078874653 | 0.9989683987 |
| **Complement.C4** | 0.0008952033478 | 0.002461451791 | 0.2166790143 | 0.9989683987 |
| **FGF.21** | -0.002600517248 | -0.002642970462 | 0.2226867873 | 0.9989683987 |
| **L.selectin** | -0.001787397862 | 0.002164534726 | 0.2414228278 | 0.9989683987 |
| **MMP.12** | 0.002845722996 | -0.0008490416694 | 0.2446503463 | 0.9989683987 |
| **RARRES2** | 0.0006662812642 | -0.0008207870631 | 0.2893759682 | 0.9989683987 |
| **FcRL2** | 0.001409784616 | -0.0003806847187 | 0.2967171324 | 0.9989683987 |
| **Myeloperoxidase** | -0.001700098875 | 0.0009619427831 | 0.3079605512 | 0.9989683987 |
| **NMNAT1** | 0.0009248453747 | -0.001046839759 | 0.3079605512 | 0.9989683987 |
| **Alpha.L.iduronidase** | -0.001586732312 | 0.0005394910755 | 0.335277798 | 0.9989683987 |
| **MMP.1.1** | 0.002617914772 | -0.0004795092825 | 0.3393039543 | 0.9989683987 |
| **Waist.Hip.Ratio** | 0.004785177115 | 0.002362838477 | 0.3474488829 | 0.9989683987 |
| **Testican.2** | 0.001418897395 | 0.0001742450372 | 0.3515676019 | 0.9989683987 |
| **PIGR** | 0.0008219686285 | -0.0002180970136 | 0.3598973853 | 0.9989683987 |
| **ESM.1** | -0.000576849828 | 0.0005601781908 | 0.4124403625 | 0.9989683987 |
| **HDL.Cholesterol** | -0.01181922425 | 0.000002421792444 | 0.4170151527 | 0.9989683987 |
| **B2.microglobulin** | 0.003914250702 | -0.0004911990806 | 0.4450856538 | 0.9989683987 |
| **CCL21** | -0.0005963455388 | -0.0004011925487 | 0.4595153953 | 0.9989683987 |
| **FAP** | -0.0001376186235 | 0.0003160726244 | 0.4595153953 | 0.9989683987 |
| **MIA** | 0.00001389760461 | -0.0009370849232 | 0.4643828978 | 0.9989683987 |
| **Interleukin.19** | -0.00005526310484 | -0.0004332867977 | 0.4742034356 | 0.9989683987 |
| **Coagulation.factor.VII** | 0.003018531584 | 0.003091150809 | 0.4791561718 | 0.9989683987 |
| **Semaphorin.3E** | -0.0002407264728 | 0.00003832858881 | 0.4791561718 | 0.9989683987 |
| **LGALS3BP** | 0.00008624623521 | 0.0003827604661 | 0.4891457976 | 0.9989683987 |
| **BMP.1** | -0.00240163659 | -0.001623841147 | 0.5043381406 | 0.9989683987 |
| **GHR** | -0.002040970852 | -0.0006354712703 | 0.5043381406 | 0.9989683987 |
| **CCL22** | 0.0004679041346 | -0.0003383886162 | 0.5197757608 | 0.9989683987 |
| **NTRK3** | -0.002482866877 | 0.0001828863074 | 0.5197757608 | 0.9989683987 |
| **TNFRSF17** | -0.0008848719987 | -0.0001668868935 | 0.5197757608 | 0.9989683987 |
| **CCL18** | -0.00118479946 | 0.001843641076 | 0.5302012868 | 0.9989683987 |
| **FCER2** | 0.0009996415821 | -0.0004107485766 | 0.5302012868 | 0.9989683987 |
| **Tryptase.beta.2** | -0.009750982018 | -0.002955007247 | 0.5302012868 | 0.9989683987 |
| **VEGFA** | 0.001761861761 | 0.001340692446 | 0.5729382548 | 0.9989683987 |
| **NRTK3** | 0.0003012171188 | 0.0001495477023 | 0.5783931804 | 0.9989683987 |

| | | | | |
|---|---|---|---|---|
| **N.CDase** | 0.002411467873 | 0.00935735488 | 0.5838724 | 0.9989683987 |
| **Osteomodulin** | 0.001871324262 | -0.00017438412 | 0.5949027425 | 0.9989683987 |
| **THBS2** | 0.0004041928107 | -0.002311829086 | 0.5949027425 | 0.9989683987 |
| **BCAM** | 0.000003496856842 | -0.00001249792955 | 0.6060272831 | 0.9989683987 |
| **IGFBP.1** | 0.002664630607 | 0.003683271563 | 0.6342370447 | 0.9989683987 |
| **SERPIN.A3** | 0.003140451133 | 0.0008758347599 | 0.6342370447 | 0.9989683987 |
| **MMP.2** | 0.000008367060625 | -0.0000792192152 | 0.6399451274 | 0.9989683987 |
| **LY9** | 0.002719537659 | 0.002161660627 | 0.6514251765 | 0.9989683987 |
| **CXCL11.soma** | 0.0007340883083 | -0.0003227192082 | 0.6571965641 | 0.9989683987 |
| **Resistin** | 0.0002396798435 | 0.001550441076 | 0.6571965641 | 0.9989683987 |
| **Body.Fat..** | -1.216026354 | -0.2231436556 | 0.6688005903 | 0.9989683987 |
| **SIGLEC1** | -0.0001177740104 | -0.00180246272 | 0.6688005903 | 0.9989683987 |
| **SKR3** | 0.00001285003897 | -0.002576840009 | 0.6746326279 | 0.9989683987 |
| **ADAMTS** | 0.003942952981 | 0.003928218319 | 0.6863552226 | 0.9989683987 |
| **NEP** | -0.003103583437 | 0.0009963272374 | 0.7040807194 | 0.9989683987 |
| **HCII** | 0.001086743038 | 0.0008045524871 | 0.7159883899 | 0.9989683987 |
| **OSM** | 0.001572286871 | -0.001098441165 | 0.7159883899 | 0.9989683987 |
| **SMPD1** | 0.002112516179 | 0.001853556137 | 0.7219684478 | 0.9989683987 |
| **ENPP7** | 0.002613330293 | 0.003970865634 | 0.7400095086 | 0.9989683987 |
| **HGFI** | 0.04255047106 | 0.04580961298 | 0.7400095086 | 0.9989683987 |
| **CD163** | 0.0009508661162 | -0.001900454481 | 0.7642869187 | 0.9989683987 |
| **CXCL9** | 0.0004576005966 | 0.0008835725259 | 0.776514776 | 0.9989683987 |
| **EZR** | 0.00005323461148 | -0.0003401201723 | 0.7826497483 | 0.9989683987 |
| **MMP.9** | -0.000681185814 | -0.0005637678863 | 0.7826497483 | 0.9989683987 |
| **GPIba** | -0.0006501529099 | -0.0009839887479 | 0.7887982685 | 0.9989683987 |
| **ICAM5** | -0.003377737211 | -0.001299728137 | 0.7887982685 | 0.9989683987 |
| **PAPP.A** | -0.001186602545 | -0.001643318436 | 0.7887982685 | 0.9989683987 |
| **Adiponectin** | -0.000368786407 | -0.003670844601 | 0.7949599725 | 0.9989683987 |
| **E.selectin** | -0.0008755952635 | -0.0007930053268 | 0.7949599725 | 0.9989683987 |
| **GDF.8** | 0.0002305888132 | 0.0002630791744 | 0.7949599725 | 0.9989683987 |
| **MRC2** | -0.0003441140358 | 0.000241747697 | 0.8073214634 | 0.9989683987 |
| **CRTAM** | -0.0003649384859 | 0.0008722371158 | 0.8135205102 | 0.9989683987 |
| **Galectin.4** | -0.0002285249335 | -0.0001310805028 | 0.8197312605 | 0.9989683987 |
| **Lymphotoxin.abeta** | -0.001000085492 | -0.004109682487 | 0.8259533386 | 0.9989683987 |
| **CHIT.1** | 0.0001064523224 | 0.001253765024 | 0.8321863666 | 0.9989683987 |
| **Contactin.4** | 0.00122158863 | 0.0008709213154 | 0.8321863666 | 0.9989683987 |
| **NCAM.120** | 0.001472576064 | -0.0002932360989 | 0.8321863666 | 0.9989683987 |

| | | | | |
|---|---|---|---|---|
| **Aminoacylase.1** | 0.001675816685 | -0.001459171278 | 0.8446837506 | 0.9989683987 |
| **Body.Mass.Index** | -0.01829311343 | -0.0128794876 | 0.8572203494 | 0.9989683987 |
| **MMP.1** | -0.0001887707847 | -0.0003103901232 | 0.8572203494 | 0.9989683987 |
| **Trypsin.2** | -0.001670578608 | -0.002872303469 | 0.8572203494 | 0.9989683987 |
| **CDL5** | -0.0005564806795 | 0.001898859675 | 0.8697930701 | 0.9989683987 |
| **Granzyme.A** | -0.0009004787999 | -0.00180643731 | 0.8697930701 | 0.9989683987 |
| **Stanniocalcin.1** | -0.002183077473 | 0.00195805378 | 0.8697930701 | 0.9989683987 |
| **TGF.alpha** | 0.0002829634784 | -0.001320731373 | 0.8697930701 | 0.9989683987 |
| **Relative.IL6.Level** | 0.0126283489 | 0.01777560992 | 0.8697930701 | 0.9989683987 |
| **TPO** | 0.000516942982 | -0.0004800328284 | 0.8887130474 | 0.9989683987 |
| **Ectodysplasin.A** | -0.0005005245063 | -0.00007858233604 | 0.9013623677 | 0.9989683987 |
| **Lysozyme.C** | -0.0002508442498 | -0.00007384676591 | 0.9013623677 | 0.9989683987 |
| **Alcohol** | -0.03447228293 | -0.08453733175 | 0.9076966385 | 0.9989683987 |
| **TNFRSF1B** | -0.0002531632742 | -0.0002662013871 | 0.9140367846 | 0.9989683987 |
| **SHBG** | 0.001450253694 | 0.0001245994045 | 0.9203824056 | 0.9989683987 |
| **CD48.antigen** | -0.0005786074031 | -0.002001623108 | 0.9267331002 | 0.9989683987 |
| **CLEC11A.e2** | -0.00523528939 | -0.003384806367 | 0.9330884662 | 0.9989683987 |
| **FCGR3A** | -0.0006931375727 | -0.00293786917 | 0.9330884662 | 0.9989683987 |
| **Granulysin** | 0.0003857477398 | 0.00001546427019 | 0.9394481002 | 0.9989683987 |
| **GZMA** | -0.0006083701915 | -0.00002283059874 | 0.9458115984 | 0.9989683987 |
| **WFIKKN2** | -0.000192622704 | 0.001309446329 | 0.9458115984 | 0.9989683987 |
| **Epigenetic.Age..Zhang.** | 0.06837951592 | 0.1544228142 | 0.952178556 | 0.9989683987 |
| **CXCL10.soma** | 0.000530649964 | -0.00005319367948 | 0.952178556 | 0.9989683987 |
| **CXCL11** | -0.001922826117 | 0.003847603819 | 0.952178556 | 0.9989683987 |
| **S100.A9** | -0.001809123652 | -0.0009185710265 | 0.952178556 | 0.9989683987 |
| **CXCL10** | -0.002239725134 | -0.002218975739 | 0.9585485674 | 0.9989683987 |
| **NOTCH1** | -0.0001156664781 | 0.0003349860589 | 0.9585485674 | 0.9989683987 |
| **VCAM1** | -0.0003345048158 | 0.00003166906158 | 0.9585485674 | 0.9989683987 |
| **Afamin** | -0.000354242315 | -0.0002201612204 | 0.9649212268 | 0.9989683987 |
| **HGF** | 0.0001379774814 | 0.0002887524682 | 0.9712961276 | 0.9989683987 |
| **CRP** | -0.001499514853 | -0.000571591246 | 0.9776728628 | 0.9989683987 |
| **EN.RAGE** | -0.001079737208 | 0.0006186750341 | 0.9776728628 | 0.9989683987 |
| **CLEC11A.e1** | -0.003310912633 | -0.002682490204 | 0.9840510252 | 0.9989683987 |
| **IGFBP.4** | -0.0001903608078 | 0.0004681879562 | 0.9840510252 | 0.9989683987 |
| **CD209.antigen** | 0.00112310137 | -0.002100367834 | 0.9904302072 | 0.9989683987 |
| **Insulin.receptor** | -0.0009746132488 | -0.0005752805747 | 1 | 1 |
